# Development of an IntelliCage based Cognitive Bias Test for Mice

**DOI:** 10.1101/2022.10.19.512853

**Authors:** Pia Kahnau, Anne Jaap, Birk Urmersbach, Kai Diederich, Lars Lewejohann

## Abstract

The cognitive bias test is used to measure the emotional state of animals with regard to future expectations. Thus, the test offers a unique possibility to assess animal welfare with regard to housing and testing conditions of laboratory animals. So far, however, the performance of such a test is time consuming and requires the presence of an experimenter. Therefore, we developed an automated and home-cage based cognitive bias test based on the IntelliCage system. We present several developmental steps to improve the experimental design leading to a successful measurement of cognitive bias in group-housed female C57BL/6J mice. The automated and home-cage based test design allows to obtain individual data from group-housed mice, to test the mice in their familiar environment, and during their active phase. By connecting the test-cage to the home-cage via a gating system, the mice participated in the test on a self-chosen schedule, indicating high motivation to actively participate in the experiment. We propose that this should have a positive effect on the animals themselves as well as on the data. Unexpectedly, the mice showed an optimistic cognitive bias after enrichment was removed and additional restraining. An optimistic expectation of the future as a consequence of worsening environmental conditions, however, can also be interpreted as an active coping strategy in which a potential profit is sought to be maximized through a higher willingness to take risks.

## Introduction

It has been shown that in both humans and animals, past experiences influences future expectations (Harding et al., 2004; Mendl et al., 2009; Paul et al., 2005). Individuals with negative experiences or in a bad state of mood are more likely to be “pessimistic” about future events and, vice versa, individuals with positive experiences or in a good mood are more likely to be “optimistic”. In the past, many tests for various species were developed to investigate the emotional state of animals (Jirkof et al., 2019). To examine the influence of emotional or affective states on expectations of future events, a number of cognitive bias tests were developed (Boleij et al., 2012; Harding et al., 2004; Hintze et al., 2018; Schlüns et al., 2017; Verbeek et al., 2014).

The common feature of these tests is the need for conditioning the subjects to scalable stimuli, e.g., odors, tones, or spatial positions. The animals learn that they will receive a reward for the stimulus at one end of the scale and a punishment for the second stimulus at the other end of the scale. After successful conditioning, the actual test follows in which ambiguous stimuli are presented to the animals. These ambiguous stimuli are located on the scale between the already known stimuli. The reaction towards these ambiguous stimuli is then measured and analyzed: It is assumed that if the response to the ambiguous stimulus is similar to the positively conditioned stimulus, the animals seem to expect a reward. They had a positive expectation of the future event, or in other words, they appear to be “optimistic”. However, if the response resembles the response of the negatively conditioned stimulus, the animals seem to have a negative expectation or seem to be “pessimistic”.

The first cognitive bias test was developed by Harding and colleagues in 2004 (Harding et al., 2004). Rats were conditioned to press a lever in response to hearing the positively associated tone-frequency to receive a reward or not to press a lever to avoid a punishment after hearing the negatively associated tone-frequency. The cognitive bias test revealed that rats kept under unpredictable housing conditions were less likely to press the lever for a reward in response to ambiguous tone-frequencies than rats kept under normal housing conditions. It was thus concluded that the negative experience rendered them ‘pessimistic’.

Although mice are the most commonly used experimental animals (Lewejohann et al., 2020), it took eight years before the first results of a cognitive bias test for mice were published (Boleij et al., 2012). Boleij and colleagues conditioned mice to various odor stimuli, which predicted either a palatable or an unpalatable food reward. First, it was shown that BALB/cJ mice were able to discriminate between odor stimuli, whereas 129P3/J mice were not. Second, it was shown that BALB/cJ mice tested under more aversive white light conditions had a higher latency in response to the ambiguous stimulus than mice tested under less aversive red-light conditions.

Further cognitive bias test methods followed in which mice were conditioned to spatial positions (Bailoo et al., 2018; Kloke et al., 2014; Novak et al., 2015; Verjat et al., 2021), to tactile stimuli (Novak et al., 2016), to different tunnel lengths (Krakenberg et al., 2019), to auditory stimuli (Jones et al., 2017), to olfactory stimuli (Resasco et al., 2021), or in an automated touchscreen-based set-up presenting different patterns on a screen (Krakenberg et al., 2019A). These studies showed that mice could be conditioned to the different stimuli and that the data plotted on the axis of stimuli increasing from negative to positive result in a sigmoidal curve (increasing s-shape slopes from the negative to the positive stimulus). These sigmoidal curves indicate that ambiguous stimuli are perceived differently compared to the conditioned stimuli, which is an important criterion for the validity of cognitive bias tests (Gygax, 2014; Hintze et al., 2018; Krakenberg et al., 2019).

So far, in all set-ups it is necessary for both the conditioning and the test itself to remove the mice from their home-cages and manually place them in the respective test set-ups. As a consequence, the animals have to be handled, taken out of their familiar environment, separated from their group members (if kept in groups) and forced to participate in the test irrespective of their current state of motivation. In fact, this may have a negative effect on the animals’ state of mind during the conditioning phase and as a result the cognitive bias test might also be influenced. This implies that in order to minimize external influence on the cognitive bias the best handling method has to be chosen (e.g., known influence of tail handling compared to cup and tunnel handling on anxiety-like behavior (Hurst & West, 2010) and that the animals have to be very well habituated to the test set-ups. Nevertheless, even with the best handling and habituation, a possibly negative influence of the separation from the home-cage and/or the group (Krohn et al., 2006; Manouze et al., 2019) as well as the experimenter’s immediate influence on the mice, and thereby the test results, must be taken into account. To overcome this shortcoming, we have developed a home-cage based cognitive bias test for mice utilizing the IntelliCage system (TSE-Systems, Germany).

The IntelliCage is a home-cage based test system that allows automated data acquisition, which can improve the reproducibility of the data (reviewed in Voikar & Gaburro, 2020). Depending on size and weight of the animals, it is possible to keep up to 16 mice in the IntelliCage as one social group. Through radio frequency identification (RFID) technology and four conditioning corners, it is possible to study activity and learning behavior in social groups (Endo et al., 2011; Kahnau et al., 2021; Krackow et al., 2010; Voikar et al., 2018).

Our test set-up consists of a home-cage, a gate (AnimalGate, TSE-Systems, Germany) and an IntelliCage (test-cage). Through the gate it is possible to separate the mice and let them individually enter the test-cage. This is especially important to allow all individuals within the group to be conditioned and tested without disturbance by group members. Another advantage is that the mice can individually decide when to enter the test-cage and participate in the experiment, rather than being coerced by an experimenter imposed schedule. As a result, the influence of the experimenter and the influence on the wake/sleep rhythm is reduced to a minimum, except for daily visual inspection and weekly cleaning of the set-up. It has already been shown that rats and mice can independently transfer themselves from their home-cages to test-cages individually to perform tasks within test-cages (Kahnau et al., 2022A; Kaupert et al., 2017; Mei et al., 2020; Rivalan et al., 2017; Winter & Schaefers, 2011). A slight disadvantage is that since only one mouse can be within the test-cage at a time and other motivated mice have to wait until this mouse has left the test-cage. However, we could show in a recent experiment with a comparable set-up that no single mouse was constantly blocking others from getting access (Kahnau et al., 2022A).

Within our automated and home-cage based test set-up, we conditioned female C57BL/6J mice to different tones. De Hoz and Nelken and Francis as well as Francis and colleagues already showed that mice were able to differentiate between different tones (De Hoz & Nelken, 2014; Francis & Kanold, 2017). Here we present our different developmental steps and results of the cognitive bias tests. Our first hypothesis was that it is possible to condition mice within the IntelliCage based set up and that the cognitive bias is influenced by the removal of enrichment and by repeated restraining. Here we present the individual developmental steps of our automated and home-cage based cognitive bias test, which were based on each other and the optimizations we implemented through previous experience. We show that it is possible to successfully condition mice in a relatively short time and measure the cognitive bias of mice, with minimal intervention and time investment by the experimenter.

## Methods

### Animals and housing conditions

In this study, three developmental steps with three different mouse groups (one developmental step per group) are presented in which different conditioning methods are described (table 1). All 36 female C57BL/6J mice were purchased from Charles River Sulzfeld, Germany. All mice were four weeks old upon arrival but were bought at different time points. Further details on the mouse groups are given at the respective developmental steps.

**Table 1:** Experimental procedure. IC = IntelliCage

|  | Group one | Group two | Group three |
| --- | --- | --- | --- |
| Developmental step | 1 | 2 | 3 |
| Year | 2020 | 2021 | 2022 |
| Conditioning protocol | Gate: passing the gate | Corner: visiting the IC corner | Corner: visiting the IC corner |
| Tone | Sequences | Frequencies | Frequencies |
| Tone length | 6.6 sec. | 0.5 and 1 sec. | 2 sec. |
| Airpuff length | 1 sec. | 1 sec. | 2 sec. |

For the establishment of the home-cage based cognitive bias test, females have been used exclusively since they can be kept in groups without complications due to little agonistic behavior. In addition, females do not show territorial behavior that excludes others (Mieske et al., 2021) and at the beginning of the development of the set-up there was a concern that individual males could occupy the gate and thus the test-cage.

We deliberately used an inbred strain to minimize genetic variability. However, despite all efforts of standardization minimal genetic drift and varying epigenetic influences can occur during breeding. In order to randomize the factors that could not be controlled for, all mice in each experiment were born and raised by different mothers and foster-mothers to ensure maximum genetic and epigenetic independence between individuals. Immediately after arrival a health inspection was performed and the mice were weighed and color-marked (edding 750, colors: black, white, red, yellow, silver) on the tail for visual identification. The mice were housed within the home-cage based set-up, no data was recorded for the first two weeks. The day after arrival, tunnel handling training to reduce handling stress (Gouveia & Hurst, 2013; Hurst & West, 2010) was started and conducted for three weeks (for a video tutorial see https://wiki.norecopa.no/index.php/Mouse_handling).

One week after arrival, all mice received RFID transponders (Euro ID, FDX-B, ISO 11784/85). The evening before the transponder transplantation, an analgesic (meloxicam 1mg/kg, Meloxidyl by CEVA) was given orally by fixing the mice in the experimenter’s hand. The transponders were implanted under isoflurane anesthesia (induction of anesthesia: 4l/min 4%; maintenance of anesthesia: 1l/min 1-2%) subcutaneously in the neck region about 1 cm behind the ears. Out of 36 mice, two mice lost their transponders until the morning after transponder implantation and the procedure had to be repeated.

One week after transponder implantation, the mice moved to the housing room where also the home-cage based experiments were conducted. The room temperature and humidity were 22°C +/- 3°C and 55% +/- 15%. The light/dark cycle was set to 12/12 hours with light off at 7 pm in winter months and at 8 pm in summer because of the switch from winter to summer time. The sunrise was simulated with a wake-up light (Philips HF 3510, 100-240 vac, 50-60 Hz, Philips Consumer Lifestyle B.V. Netherlands) half an hour before the room-light switched on. The wake-up light was placed on the ground in a corner of the housing room with the light directed towards the animals. The light intensity increased gradually and reached the full intensity at 7/8 am (depending on season). The daily visual health inspection was performed between 7/8 am to 10 am (depending on season). The home-cage set-up was cleaned once a week. Bedding, nesting material, and enrichment items were replaced. A small handful of old bedding was transferred to the new home-cage. On the same day, the mice were weighed and re-color-marked.

### Home-cage based set-up

In all developmental steps, the same home-cage based set-up was used. This set-up (figure 1) consisted of three compartments: a home-cage, a gate (AnimalGate), and a test-cage (IntelliCage). As the gate had doors and infrared barriers, it was possible to allow only one mouse at a time to pass through the gate from the home-cage into the IntelliCage (IC, test-cage). All other mice of the social group had to wait until the one mouse within the IC moved back through the gate into the home-cage.

**Figure 1:**
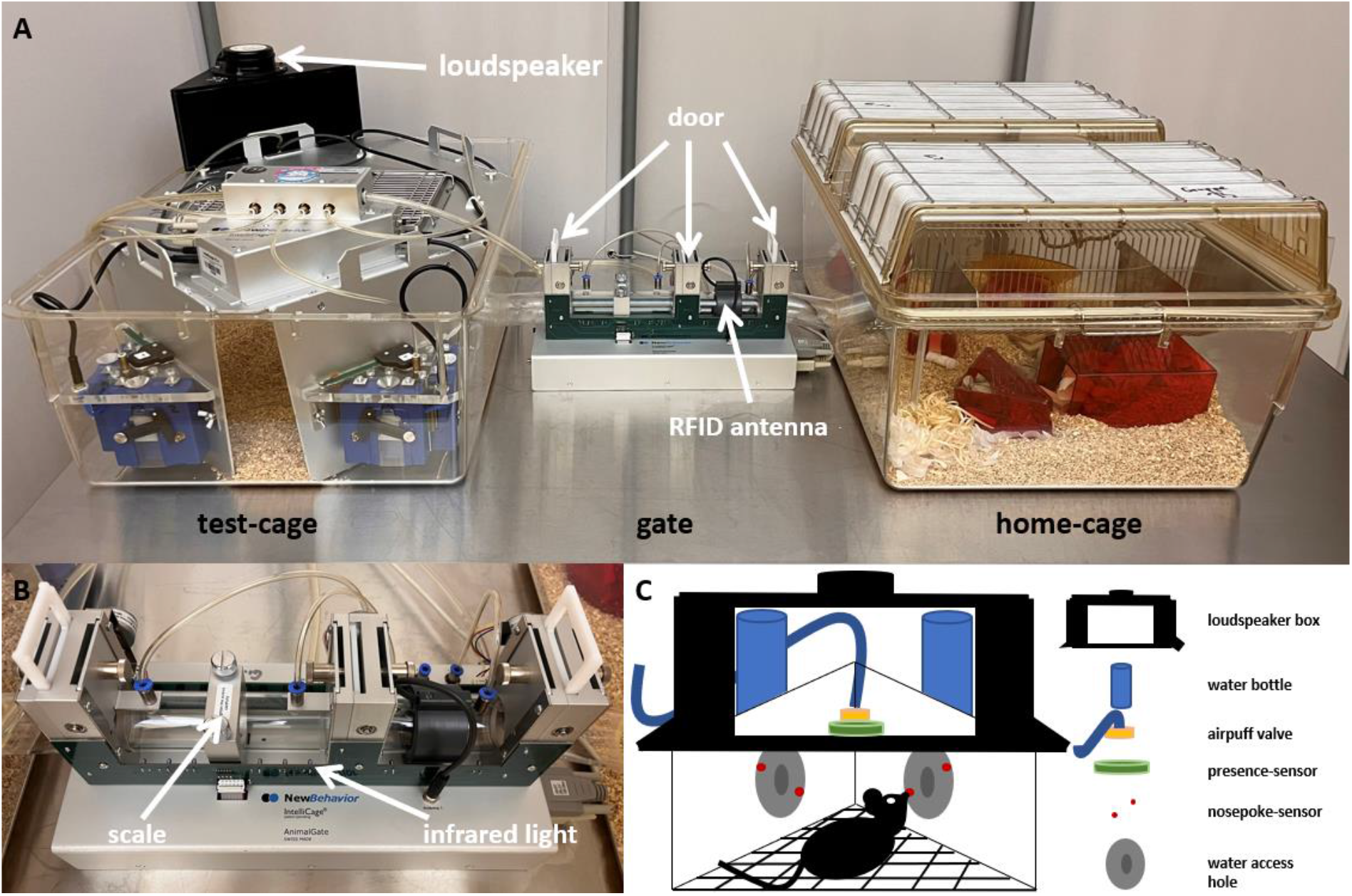
Home-cage based set-up based on the IntelliCage system. A: The set-up consisted of the IntelliCage used as the test-cage, which is connected through the AnimalGate to the home-cage. The IntelliCage was equipped with four conditioning corners and bedding. The home-cage was equipped with bedding, nesting, enrichment and food *ad libitum* (not shown here). The AnimalGate had three doors, one radio frequency identification (RFID) antenna. B: In addition, the AnimalGate had eight infrared barriers and one scale to measure the animal’s weight during each gate passage. C: Within the IntelliCage corners, water could be provided. In addition, each corner had one radio frequency identification antenna, one presence-sensor, one airpuff-valve, two water dispensers and two doors.

The home-cage was a Makrolon type IV cage (floor space 2065 cm^2^) with a filtertop equipped with 3-4 cm bedding (Poplar Granulate 2-3 mm, Altromin, Germany), two red triangle-shaped houses (“TheMouseHouse”, Tecniplast, Italy), nesting material (eight papers, six paper nesting stripes and six cotton rolls), four wooden bars to chew on, and food *ad libitum* (autoclaved pellet diet, LAS QCDiet, Rod 16, Lasvendi, Germany). Within the home-cage was also an acrylic tube (4 cm diameter, 17.5 cm long), which was used for tunnel handling to reduce handling stress. Mouse group three additionally received nesting materials upon weekly changing: folded paper stripes, mid coarse wood wool and square hemp pads. Also one resting platform and a running disk (InnoDome with InnoWheel, Bio-Serv) was placed within the home-cage and the mice received weekly changing toys filled with millet (organic peeled golden millet, Bohlsener Mühle) once per week.

The IC is a computer and RFID technology based test system with four conditioning corners. Each corner contained an RFID antenna at the corner entrance, a presence-sensor, which detected differences in temperature, two nosepoke-infrared-sensors, two doors through which the water access can be regulated, two water dispensers, and an airpuff valve for the possibility of a mild punishment (0.5 bar). Depending on the conditioning method, one or four IC corners were active, in which water was provided. The IC contained only bedding material.

In order to perform experiments within the IC system, it is necessary to habituate the mice to the system first. The mice had to learn how to pass through the AnimalGate and where to access water within the IC. For this purpose, the mice were habituated gradually to the AnimalGate and IC doors. Initially, all AnimalGate and IC doors were permanently open (phase: ‘all doors open’). Thus, it was possible for all mice to move freely within the system. As a next step, the doors of the AnimalGate were closed, and opened only when a mouse entered the AnimalGate, which is similar to the next IC habituation step when the corner doors were closed and opened due to a visit (phase: ‘visit open doors’). In the final phase of habituation, only one mouse could stay in the IC, and the IC doors could only be opened with a nosepoke.

### Conditioning protocol

The basic requirement for performing a cognitive bias test is to condition the animals to scalable stimuli. In our study, the mice were conditioned to auditory stimuli. Three different conditioning protocols were performed with each of the different mouse groups. Common to all protocols was that the mice had to learn that for one presented tone (positive tone); they received water as a reward, if they made a nosepoke within the IC corner (correct behavior). For another tone (negative tone), they received an airpuff as a punishment, if they made a nosepoke (incorrect behavior). If the mice did not make a nosepoke after hearing the positive tone (incorrect behavior), they received no water. If the mice did not make a nosepoke after hearing the negative tone (correct behavior), they did not receive an airpuff (table 2). All tones were created by using the online tool onlinetonegenerator.com and Audacity (AudacityCross-Platform Sound Editor).

**Table 2:** Description of the possible events during the conditioning within the IntelliCage corner.

| Nosepoke | Positive Tone | Negative Tone |
| --- | --- | --- |
| Yes | water | airpuff |
| No | nothing | nothing |

Since the mice only had the opportunity to drink water in the IC, it was necessary to monitor whether all mice drank daily. If a mouse did not drink for 24 h, the mouse was offered water in a separate cage for 15 minutes. After these 15 minutes, they were placed back in the home-cage. If drinking did not occur in the IC for three consecutive days, these mice were taken out of the experiment by allowing them access to water within the IC corner without tones. These mice were no longer participating in the conditioning phase and cognitive bias test, but were still left in the group, leaving the social structure unchanged throughout the experiment.

For more clarity, the individual development steps are described individually below. The respective results and conclusions follow the method description of the individual development steps.

### Analysis

Data analysis was done with the open-source statistical software R (version 4.0.3, RCoreTeam, 2020). For data visualization the R package ggplot2 (Wickham, 2016) was used. Model assumptions were inspected visually first by Q-Q plots, secondly by visualizing variance homogeneity of the residuals versus fitted values.

#### Analysis of data from gate conditioning protocol

For the gate conditioning protocol (detailed description below), the mice first had to learn which corner was the active corner. Therefore, the visit number of the active corner was compared to the visit number of the inactive corners for each mouse per day during the first 14 days (when only the positive tone was presented). A visit was recorded by the IC-system each time a mouse entered a corner and both the RFID transponder number was detected and the presence-sensor was activated. The visit number was used as the outcome in a linear mixed-effects model (R package nlme (Pinheiro et al., 2020)). The experimental days were used as a fixed effect (factor with 14 levels). The type of visit (factor with two levels: visits in active corners versus visits in inactive corners) and the interaction of type of visits and day were used also as fixed effects. The variable ‘days nested in animals’ were set as a random effect. Sum-contrasts were used for days and type of visits.

For the evaluation of the two gate conditioning runs, the frequency with which the mice passed the AnimalGate was first determined for each mouse for each day, i.e., how often mice were presented with tone-sequences. The duration from entering to leaving the IC is defined as IC-session. From this, it was determined how often the positive and negative tone-sequences were played (per animal, per day). Next, it was determined how often the mice visited the active corner and made nosepokes on the nosepoke-sensor during the positive and negative tone-sequence IC-sessions. The number of nosepokes was used as the outcome in a linear mixed-effects model (R package nlme). In this model, the experimental days were defined as days and used as a fixed effect (factor with nine levels in AnimalGate conditioning run 1, factor with 14 levels in AnimalGate conditioning run 2). Within the statistical model, the type of tone-sequence (two level factor: positive versus negative tone-sequence) and the interaction of type of tone-sequence and day were also used as fixed effects. Sum-contrasts were used for day and type of tone-sequence. The variable ‘experimental days nested in animals’ was set as a random effect.

#### Analysis of data from corner conditioning protocol

For the evaluation of the corner conditioning protocol (detailed description below), the frequency with which the mice visited the active corner within the IC was first determined for each mouse for each day, i.e., how often mice were presented with tone-frequencies (inactive corners were blocked with a plug). From this, it was determined how often the positive and negative tones were played (per animal, per day). Next, it was determined how often the mice visited the active corner and made nosepokes at the nosepoke-sensor during the positive and negative tone. The number of nosepokes was used as the outcome in a linear mixed-effects model (R package nlme). In this model, the experimental days were defined as days and used as a fixed effect (factor with 48 levels). Within the statistical model, the type of tone-frequency (two level factor: positive tone-frequency versus negative tone-frequency) and the interaction of type of tone-frequency and day were also used as fixed effects. Sum-contrasts were used for day and type of tone-frequency. The variable ‘experimental days nested in animals’ was set as a random effect. To test for effects of interaction of day and tone-frequency, *post hoc* comparison was conducted (R package emmeans (Lenth, 2020)).

#### Learning success for visit conditioning

Descriptive statistics were used to assess individual learning success by observing correct nosepoke behavior. Correct nosepoke behavior at the positive tone is defined as a corner visit during which at least one nosepoke was made. Correct nosepoke behavior for the negative tone is defined as a corner visit without a nosepoke. For each mouse, we first determined how many positive tone trials and negative tone trials had occurred. Then, the numbers of positive tone trials with nosepokes and the number of negative tone trials without nosepokes were determined. Since the probabilities for the positive and negative tone trials were different, percentage values were calculated. From this, the corrected nosepoke behavior was plotted for each animal individually. The learning criterion was set as follows: First, it was checked whether the values for the positive and negative tone are above the 50% chance level. Then, on 75% of the conditioning days, the correct nosepoke behavior had to be above the chance level in order to reach the learning criterion.

#### Cognitive bias test

For the cognitive bias test, the mice were presented with three additional (ambiguous) tones. First, for each mouse it was determined how many nosepokes they made in response to the five different tones. The number of nosepokes was used as the outcome in a linear mixed-effects model (R package nlme). In this model, the tones (factor with five levels) and measurement (cognitive bias test 1: factor with three levels (baseline measurement 1, negative conditions and baseline measurement 2), cognitive bias test 2: factor with four levels (baseline measurement 1 and 2, negative conditions and baseline measurement 3) and the interaction were used as fixed effects. The variabe ‘treatment nested in animals’ was set as a random effect. If the model indicated a significant effect of treatment or tone, we conducted a pairwise post hoc analysis (R package emmeans).

#### Body weight and IntelliCage behavior

For the evaluation of body weight, number of nosepokes and visits, the corresponding values were determined for each animal for each day. These three variables were used as the outcome in three different linear mixed-effects models (R package nlme). Treatment (group two: factor with eight levels (0%, 5%, 10%, 16%, 20%, 33% and 50% probability of negative tone and visit open doors, group three: factor with seven levels (0%, 20% and 50% probability of negative tone, nosepoke open doors, baseline measurement and negative conditions), day (group two: factor with 75 levels, group three: factor with 100 levels) and the interaction of treatment and day was used as a fixed effect. The variable ‘experimental days nested in animals’ were set as a random effect.

## Developmental Step 1

### Methods

#### Animals

The twelve female mice of group one arrived at the institute in February 2019. At the start of the first developmental step, the mice were seven weeks old. After the experiment presented here, the mice were 18 weeks old and used in home-cage based learning tasks (data not published) and in a consumer demand test, which was also performed within the home-cage based set-up presented here (Kahnau et al., 2022A). The mice started barbering behavior at the age of 18 weeks and immediately following the experiment presented here. Barbering behavior is commonly found in C57BL/6J mice (Kahnau et al., 2022B; Sarna et al., 2000). The reason for this behavior is not yet understood.

#### Gate conditioning protocol

The gate conditioning protocol was pre-registered in Animal Study Registry (animalstudyregistry.org, doi: 10.17590/asr.0000121). The mice were conditioned to tone-sequences. These sequences had a play time of 6.6 seconds at a frequency of 8 kHz and comprised either short tone-sequences with long breaks or long tone-sequences with short breaks (figure 2).

**Figure 2:**
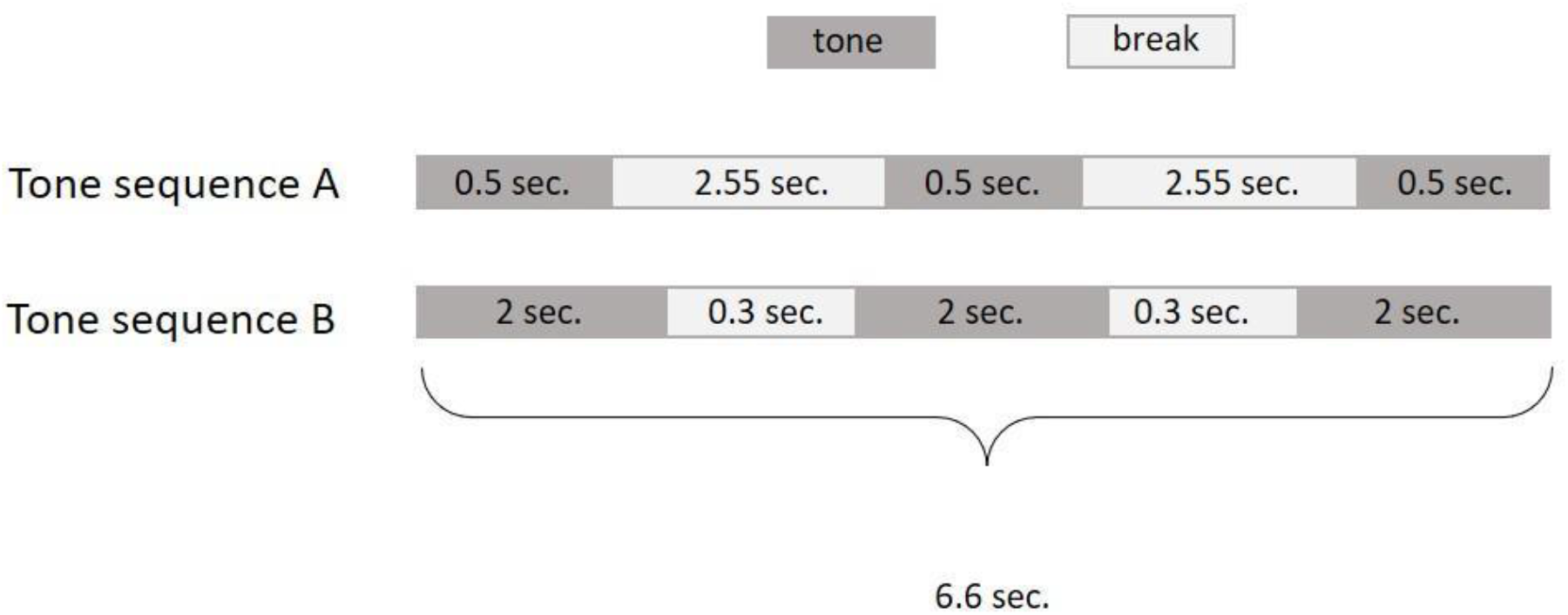
Tone-sequences used for AnimalGate conditioning.

Each mouse was randomly assigned one of two tone-sequences, thus six out of twelve mice had tone-sequence A and the other six had tone-sequence B as the positive tone stimulus. The other tone-sequence was consequently the negative stimulus. One loudspeaker was placed on top of the IC (on the grid) facing in the direction of the IC inside, allowing the mice to hear the tone-sequences (see loudspeaker position 1 in figure 1). The tone-sequences were played when entering the IC after passing through the gate.

Within the IC, each mouse was randomly assigned one active corner (three mice per corner), in which the mice received either the water reward or an airpuff punishment depending on the tone-sequence. Visiting the other three corners had no consequences.

The mice had to learn first which corner their active corner was (one out of four) and second that a tone was played every time they entered the IC through the gate. This corner and positive tone conditioning ran for 14 days. When visiting the active corner and activating the nosepoke-sensor, the doors were opened for five seconds. To prevent the mice from staying too long inside the corner, an airpuff was released after another five seconds. To open the doors within the IC corner again, the IC had to be left through the gate (end of IC-Session). By re-entering the IC, a new trial was initiated.

After corner and positive tone conditioning, the negative tone-sequence was added. To prevent the mice from having too many negative experiences directly at the beginning of the conditioning phase, the probability of the negative tone being played was increased successively. Therefore, two runs were carried out. For gate conditioning run 1, the probability of playing the negative tone was 33%. For gate conditioning run 2, the probability of playing the negative tone was 50%. To initiate a new trial, the IC had to be re-entered through the gate, i.e., mice that could not drink after a negative tone-sequence or did not drink after a positive tone-sequence had to leave and re-enter the IC for the next chance to drink.

### Results

#### Corner and positive tone-sequence conditioning

The mice first had to learn which corner was the assigned active corner. Over a period of 14 days, the animals were successfully conditioned to the active corner. (main effect visits: *F*_1,154_ = 225.44, p < 0.0001). The overall number of visits decreased over the experimental days (interaction: *F*_13,154_ = 6.63, p < 0.0001, figure 3).

**Figure 3:**
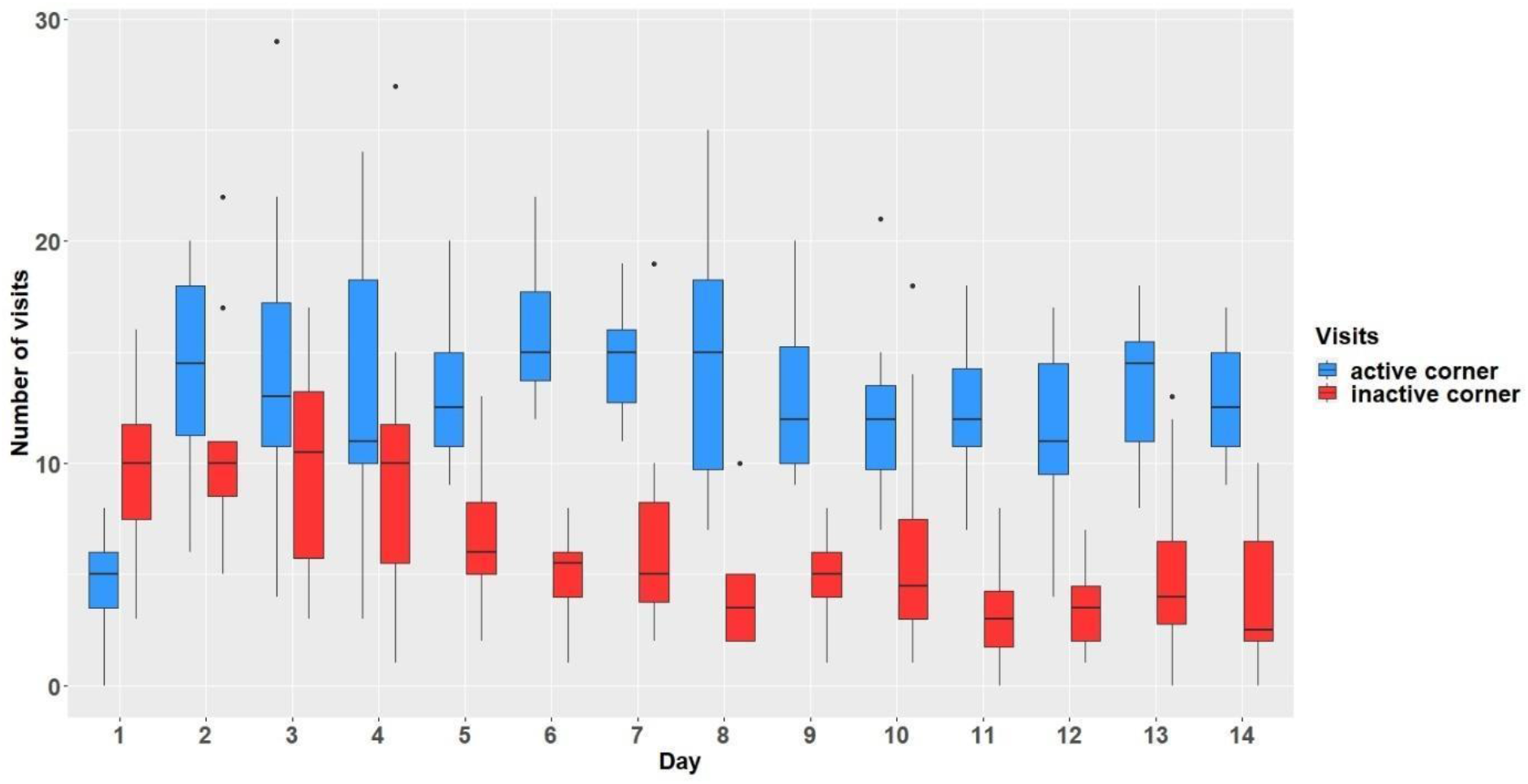
Comparison of visit numbers in active and inactive corners. The y-axis shows the number of visits which were made within the active and inactive corners. The x-axis shows the experimental days.

#### Gate conditioning protocol

The mice had to learn to make nosepokes after hearing positive tone-sequences and refrain from making nosepokes after hearing negative tone-sequences. In gate conditioning run 1 with 33% chance of hearing a negative tone sequence (figure 4), the mice did not make more or less nosepokes after hearing positive or negative tone-sequences on average (main effect tone-sequence: F_1,90_ = 0.22; p = 0.64). The mice did not learn to differentiate between tone-sequences over time (interaction: F_8,90_ = 0.82; p = 0.59). However, the mice made fewer nosepokes regardless of tone-sequences over time (main effect day: F_8,80_ = 4.58; p = 0.0001). In gate conditioning run 2 with the chance of hearing a negative tone sequence reduced to 50% (figure 5), the mice made, on average, more nosepokes for the positive tone-sequence (main effect tone-sequence: F_1,77_ = 18.9; p < 0.0001) but did not learn to differentiate between the tone-sequences (interaction: F_6,77_ = 0.62; p = 0.71). During run 2 the mice made more nosepokes over time regardless of tone-sequences (main effect day: F_6,66_ = 2.45; p = 0.03).

**Figure 4:**
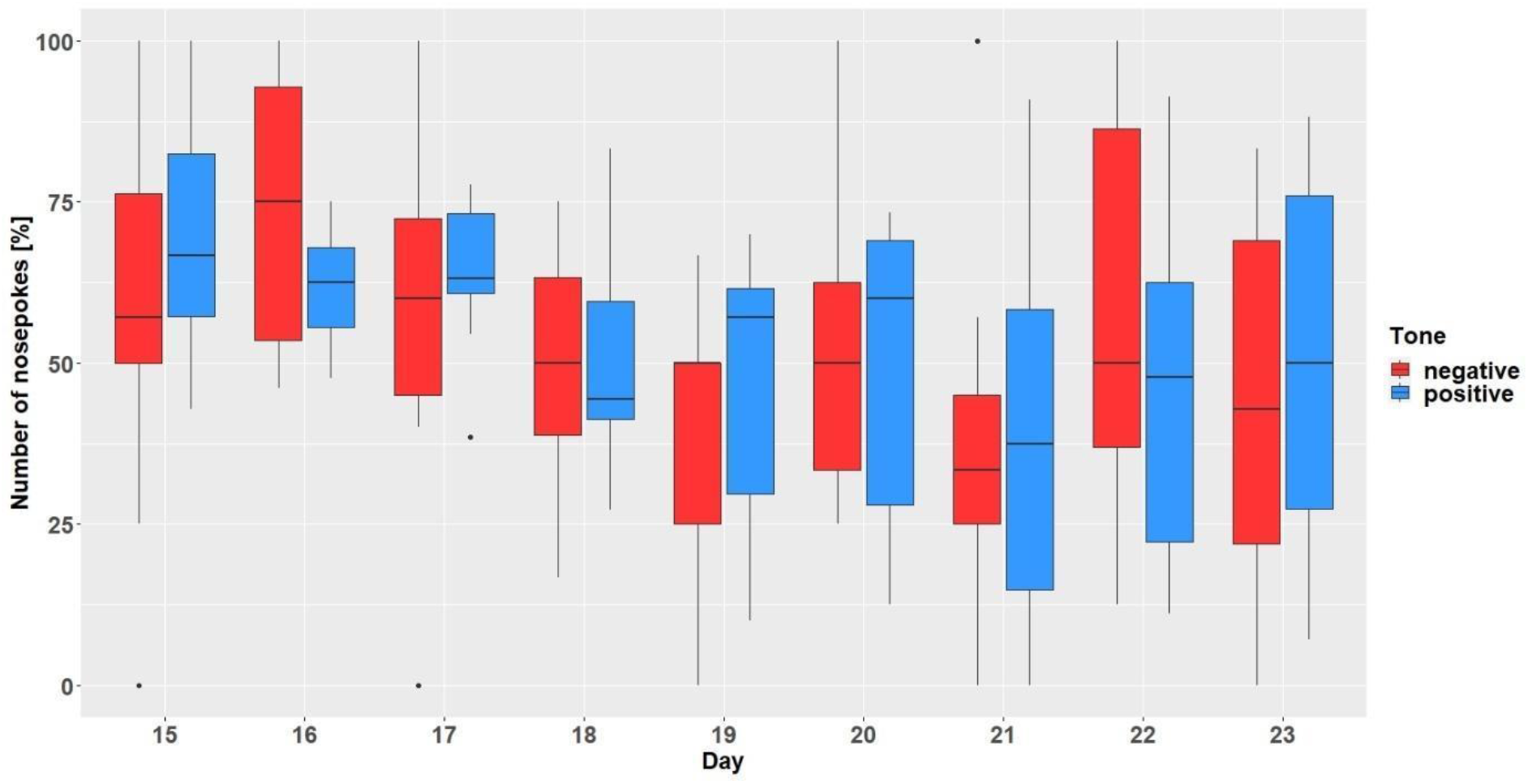
Gate conditioning run 1. The y-axis shows the number of nosepokes which were made in response to the presented tone-frequencies. Number of nosepokes are given in percent since the probability of the two tone-frequencies being played was different (positive = 67%, negative = 33%). After hearing a positive tone-frequency, a nosepoke had to be made, but not after hearing a negative tone-frequency. The x-axis shows the experimental days.

**Figure 5:**
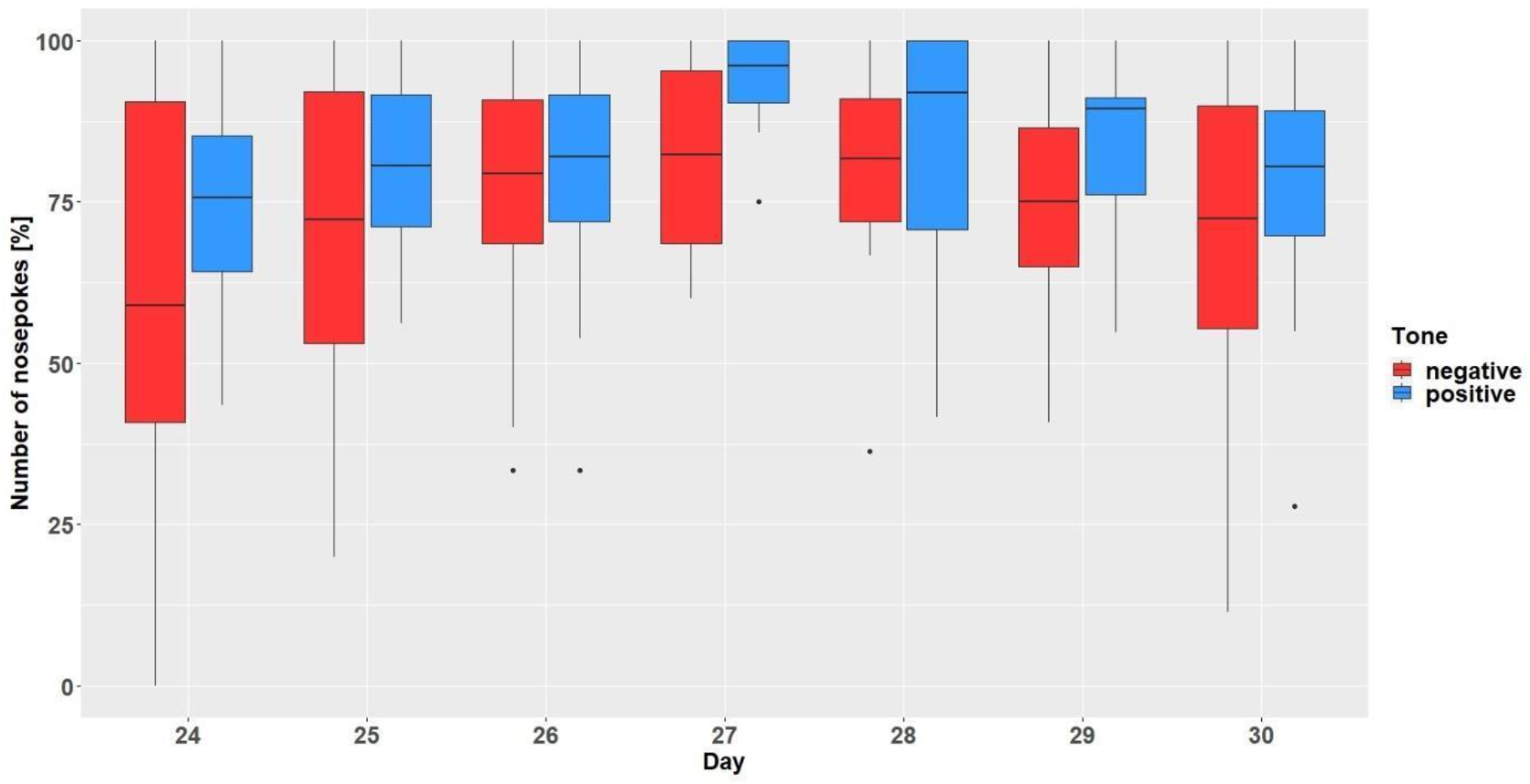
Gate conditioning run 2. The y-axis shows the number of nosepokes which were made in response to the presented tone-frequencies. The probability of the two tone-frequencies being played was 50:50. After hearing a positive tone-frequency, a nosepoke had to be made, but not after hearing a negative tone-frequency. The x-axis shows the experimental days.

### Discussion

The first developmental step was described as ‘gate conditioning protocol’, where tone-sequences were played whenever a mouse passed the gate and entered the IC. The initial idea of using tone-sequences was to easily create ambiguous sequences once the positive and negative sequences were successfully conditioned. Although it was possible to condition the mice to their respective randomly assigned IC corner, the mice were not able to distinguish between two tone-sequences. The mice were unable to associate a water reward with one tone-sequence and a mild airpuff punishment with another tone-sequence. The unsuccessful conditioning could have different reasons.

First, mouse-specific ultrasonic vocalization series can have a length of two seconds. They are variable in their sequence but are released at a more or less constant frequency. There are also short sequences (a few milliseconds long) that vary in both sequence and frequency (Ehret, 2018). Our artificially created, very static tone sequences at constant frequency have a length of 6.6 seconds, which may be too long to be perceived as relevant for the mice. The tone-sequences might have shown better results if shortened. To the best of our knowledge, there have been no experiments to condition mice to artificially created tone-sequences like the ones we used during developmental step 1. However, past studies showed the possibility to condition mice to tones, namely tone-frequencies (De Hoz & Nelken, 2014; Jones et al., 2017). Therefore, we decided to use tone-frequencies instead for the next developmental step. Second, the timing of when the tone-sequences during gate conditioning were presented was not optimal. Tones were initiated by each pass through the gate and played when the IC was entered. Whether the mouse then also directly visited the IC corner was probably dependent on how strong the motivation to drink was. Thereby it might be possible that too much time passed between the tone and the actual corner visit, and thus, no association was established between these two events. The timing between stimulus presentation and event onset is important for successful conditioning, as shown, for example, by clicker training (Lattal, 2010). Therefore, we decided to change the time point of tone presentation and relocated the conditioning completely to the IC corner. From then on, the sound was played when the mouse entered the IC corner. This improvement reduced the time span from the presentation of the stimulus to the corresponding nosepoke behavior to a minimum. To prevent a possible overlap effect of the unsuccessful conditioning on the next developmental step, we continued to work with a naïve mouse group.

## Developmental Step 2

### Methods

#### Animals

The twelve female mice of group two arrived at the institute in October 2019. At the start of the second developmental step presented here, the mice were 14 weeks old. The mice started barbering behavior at an age of 20 weeks, during the conditioning phase. At the end of the experiment, the mice were 26 weeks old used in another experiment to develop a conditioned place preference test to assess severity of experimental procedures (publication in preparation).

#### Corner conditioning protocol

Since the gate conditioning protocol was not successful in group one, we improved the conditioning protocol and decided to condition no longer to tone-sequences but to tone-frequencies.

The hearing range of mice is between 2 kHz and 70 kHz (Heffner & Heffner, 2007). To find different frequencies with equal sound pressure levels (SPL) in the corner, a measuring microphone (miniDSP Umik-1 calibrated USB microphone) and the software Room EQ Wizard (https://www.roomeqwizard.com) were used. In a study of de Hoz and Nelken, mice were successfully conditioned to tone-frequencies between 6 kHz and 13 kHz (De Hoz & Nelken, 2014). The same frequency range was used for our study. With a digital signal processor (miniDSP 2×4 https://www.minidsp.com), the SPL of the played tone was optimized, to ensure that all tones were played at the same volume within the corner. This was done to ensure that variations in SPL stemming from the speaker and confined space in which they are played were as small as possible. However, it must be emphasized that the perception of mice differs from that of humans and that there possibly are influences, which we are unable to detect and/or assess. In addition to different tones, we decided to change the time when the tones were played. For the corner conditioning protocol, tone-frequencies (positive or negative tone) were played when entering an IC corner instead of when leaving the gate and entering the IC. One single corner within the IC was chosen as the active corner for all mice to set the focus of the mice to this corner and to ensure that the tone quality was the same for all mice. All other corners were made unreachable by 3D printed plugs made from gray polylactic acid (PLA). In order to initiate a new trial, the mice had to re-enter the active corner. During one IC-session, multiple trials could be initiated by the mouse re-entering the active corner without having to leave the IC again (as it was the case for gate conditioning protocol). Within the active corner and after hearing the positive tone-frequency, the IC doors could be opened by a nosepoke for seven seconds.

To play the tone-frequencies, one loudspeaker was placed on top of the active corner directed towards the inside of the corner, so the mice were able to hear the tones. In order to be able to position the loudspeaker, it was integrated into a black 3D printed box (figure 1 C).

The tone-frequency at one end of the scale was 6.814 kHz at 70 decibel (dB), the other tone-frequency on the other end of the scale was 13.629 kHz at 70 dB. At the beginning of the conditioning phase, only the positive tone-frequency was played during a visit in the active IC corner. The probability of the negative tone-frequency was increased progressively to avoid too many negative experiences at the beginning of the experiment (supplementary material table S1).

The tone-frequencies had at first a length of 0.5 seconds. The tone length was extended to one second on experimental day 20. During the experimental phase, there were several technical problems and therefore, data for some days were lost. On several occasions, the body weight of the animals could not be recorded due to the AnimalGate being blocked by bedding material. Removing the bedding from the AnimalGate solved this problem. An unexpected failure of the control unit led to missing data recording on days 23, 99, 104, 105. The whole IC system had to be restarted to resolve these failures.

### Results

#### Corner conditioning protocol

After visiting the active corner, one out of two tone-frequencies was randomly presented. In total (figure 6), the mice made more nosepokes at the positive tone compared to the number of nosepokes made at the negative tone-frequency (main effect tone: *F*_1,452_ = 795, p < 0.0001). The mice differentiated between the two tone frequencies after the tone length was increased to 1 second on experimental day 20 (interaction: *F*_47,452_ = 9.51, p < 0.0001, table S3 in supplements). In addition, the mice made less nosepokes in total after day 20 (main effect experimental day: *F*_47,473_ = 5.39, p < 0.0001).

**Figure 6:**
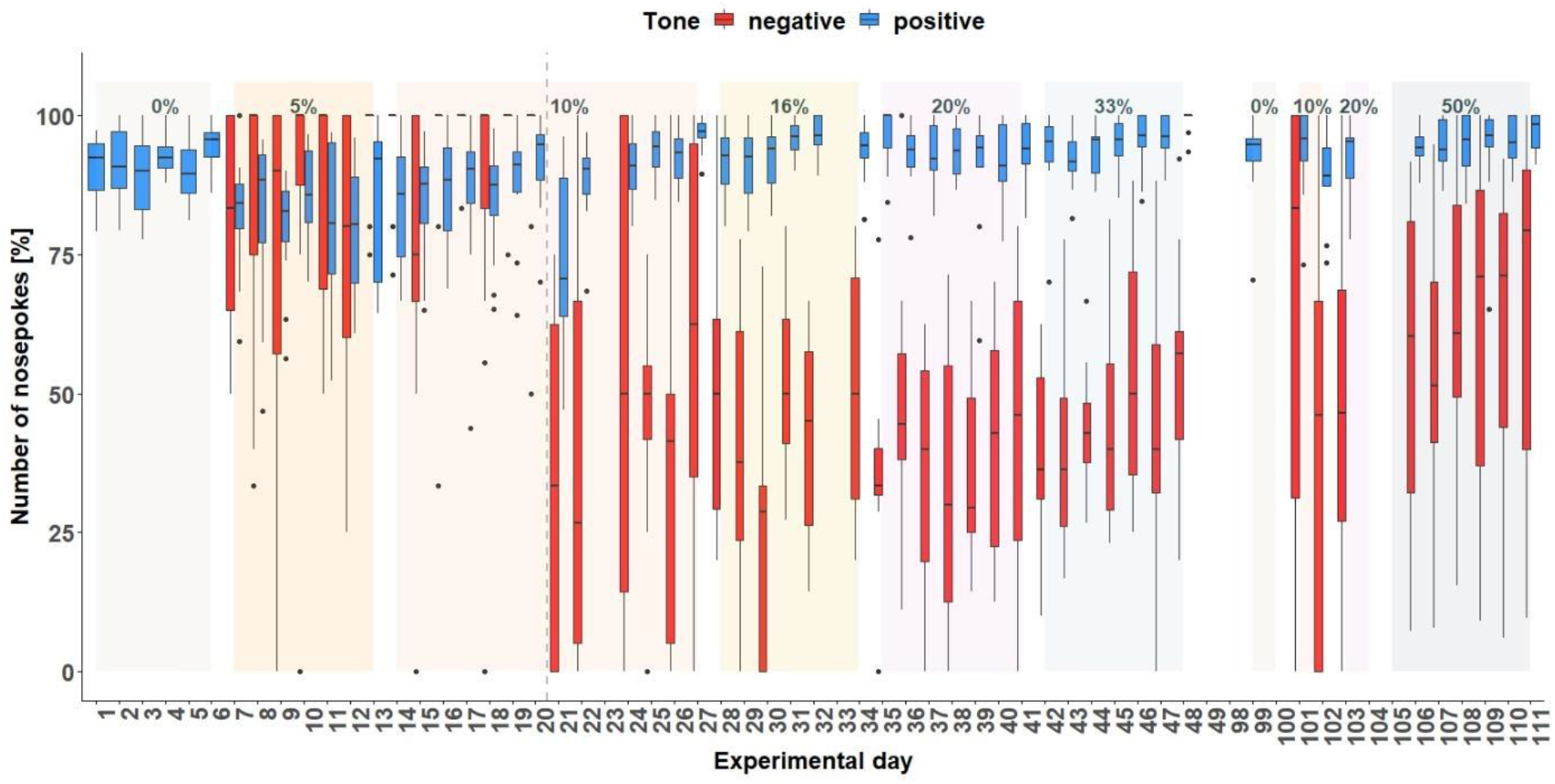
Corner conditioning group two. Number of nosepokes in percent made in response to two different tone-frequencies. The data for experimental day 23, 99, 104, 105 is missing due to technical problems with the IntelliCage system. No data from day 49 to 84 is available, because the mice were not in the home-cage based set-up as the set-up had to be maintained. From experimental day 85 the mice were kept in the set-up again. In order to habituate the mice to the set-up again, no sounds were played on days 85 to 98. On the y-axis, the number of nosepokes in percent is shown. The x-axis shows the experimental days. The dashed line marks the time point when the tone length was increased to one second. 0% = no negative tone, 5% = 5% negative tone probability, 10% = 10 percent negative tone probability, 16% = 16% negative tone probability, 20% = 20% negative tone probability, 33% = 33% negative tone probability, 50% = 50% negative tone probability

#### Individual learning success

Since the results are considered for each mouse, the results are evaluated descriptively. The individual learning success is considered during the time period when the negative tone was played with a probability of 33% (figure 7) and 50% (figure 8). These were chosen because the negative tone was played enough times to allow a meaningful comparison of the nosepoke behavior.

**Figure 7:**
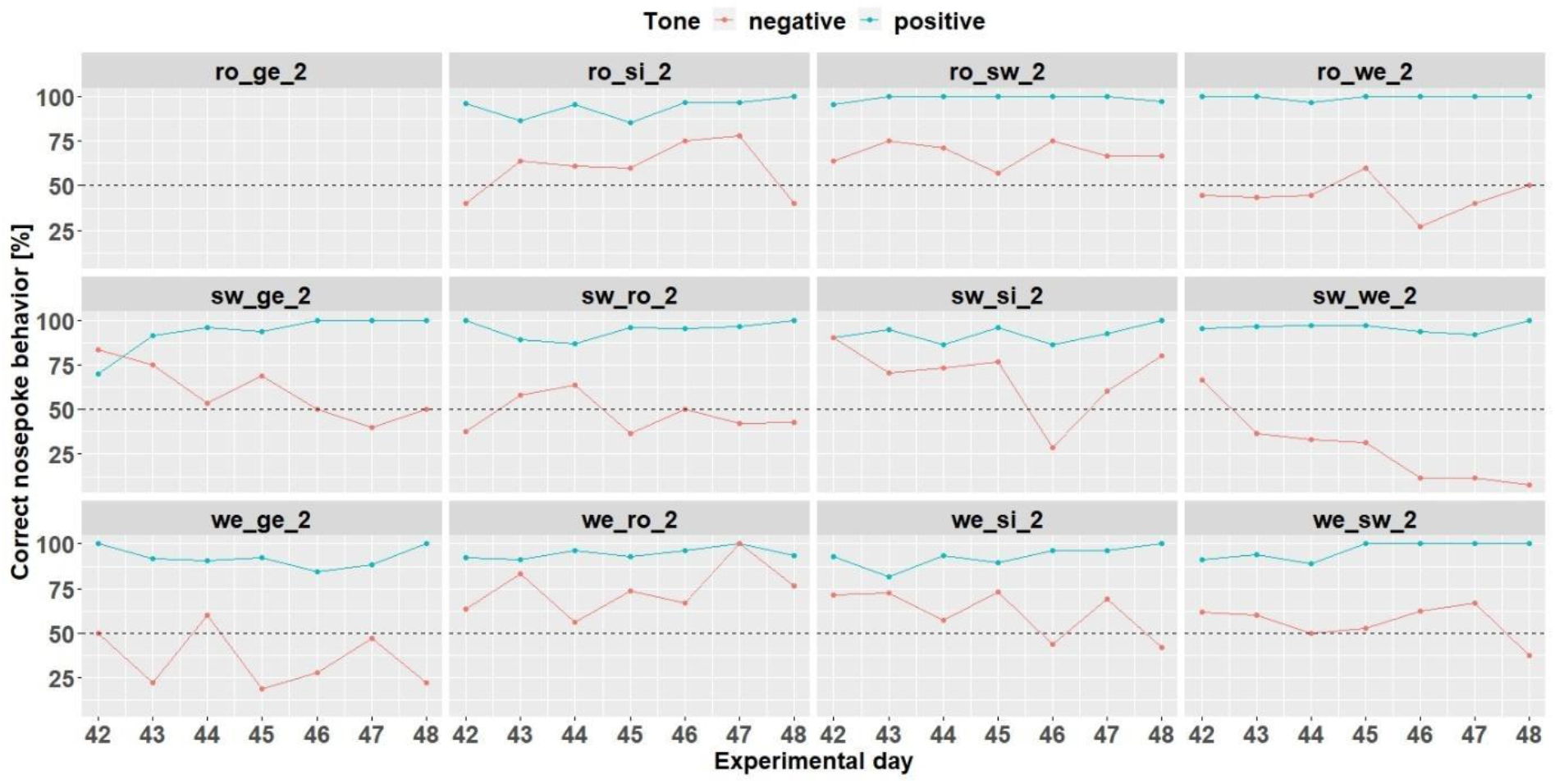
Individual learning success during conditioning when the negative tone-frequency was presented with a probability of 33%. Mouse ro_ge_2 was taken out of the experiment. Learning criterion 75% of correct nosepoke behavior over 50%.

**Figure 8:**
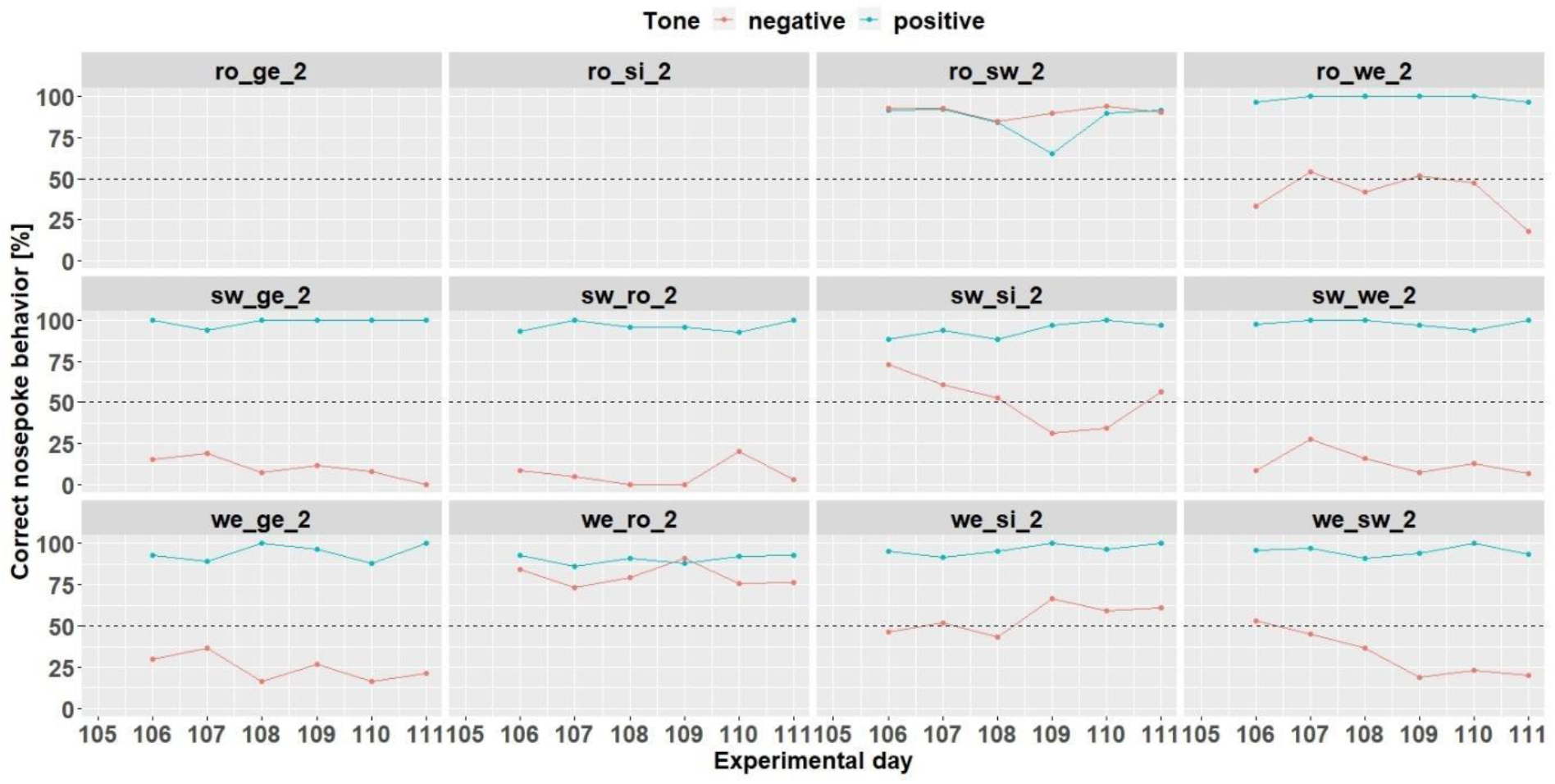
Individual learning success during conditioning when the negative tone-frequency was presented with a probability of 50%. The data of day 105 is missing due to technical problems with the IntelliCage system. Ro_ge_2 and ro_si_2 did not participate any longer in the experiment. Learning criterion 9 trials over 50% out of 12.

At the time when the negative tone was played with a probability of 33%, seven mice (ro_si_2, ro sw 2, sw ge 2, sw si 2, we ro 2, we si 2, and we sw 2) out of 12 mice reached the learning criterion. Mouse ro_ge-2 stopped to drink before the negative tone was played with a probability of 33%.

Increasing the probability of the negative tone to 50% resulted in more incorrect nosepoke behavior in response to the negative tone. Only four mice (ro_sw_2, sw_si_2, we_ro_2 and we_si_2) out of 12 mice reached the learning criterion (75% of correct nosepoke behavior over 50%). The mice ro_ge_2 and ro_si_2 did not drink and were taken out of the experiment.

#### Body weight and IntelliCage behavior

Body weight, number of licks, and number of visits were recorded throughout the experimental period. (figure 9). Body weight was influenced by the treatment (*F*_7,803_ = 2.33, p = 0.02) as well as by the experimental day (*F*_1,803_ = 211, p < 0.0001). Also the interaction treatment and day had an influence on body weight (*F*_7,803_ = 2,36, p = 0.02). Also, the number of licks over time was influenced by treatment (*F*_7,813_ = 20.71, p < 0.0001) as well as experimental day (*F*_1,813_ = 14,3, p > 0.0001). This influence seems to be particularly strong on individual experimental days (interaction: *F*_17,813_ = 7,99, p < 0.0001), which is also reflected in the number of visits (interaction: *F*_7,813_ = 22.31, p < 0.0001). These were also influenced by the treatment (*F*_7,814_ = 47, p < 0.0001) but not influenced by the experimental day (*F*_1,815_ = 0.05, p = 0.83).

**Figure 9:**
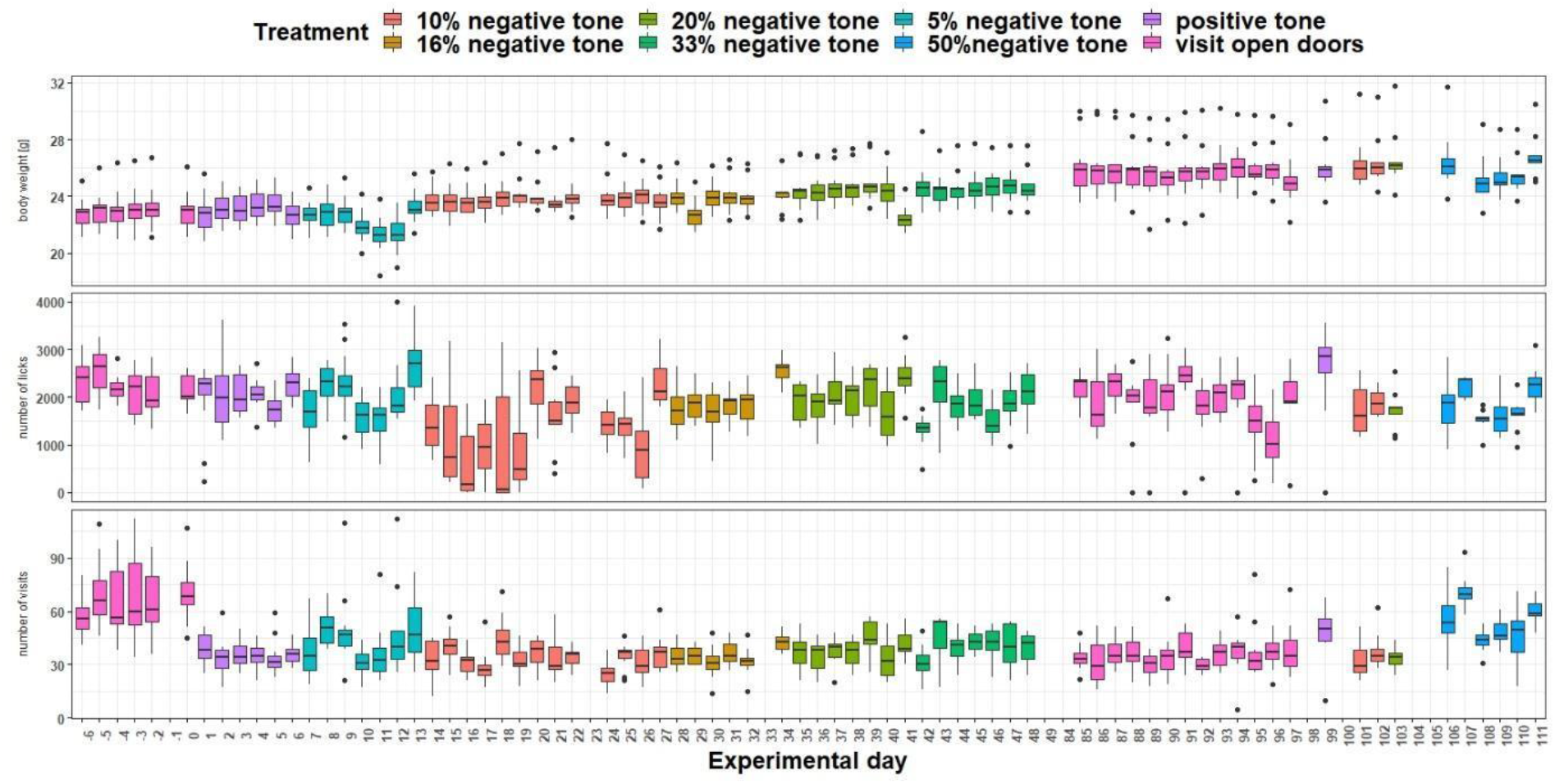
Measurement of body weight, IntelliCage corner visits and lick number over time. The x-axis shows the experimental days. On the y-axis first the body weight, second the lick number and third the visit number is shown. Different tones with different playback probabilities were presented throughout the experimental period (treatment). The data of experimental day −1, 23, 33, 98, 100, 104 and 105 are missing, due to technical issues. During experimental days 49 to 84 no tones were played.

### Discussion

The second developmental step was described as ‘corner conditioning protocol’, where tone-sequences were played whenever a mouse visited the active IC corner. With this protocol, it was possible for the first time for single mice to distinguish between two different tone-frequencies within the set-up presented here. Two mice ceased drinking in the IC during the conditioning phase. Therefore, these mice were excluded from the experiment, i.e., for them the tone presentation was turned off and they were able to open the doors by a nosepoke at each visit. Dropouts also occurred in other studies, where individual animals did not reach the learning criterion and thus the actual test phase (e.g., Bračić et al., 2022; Hintze et al., 2018; Kloke et al., 2014; Krakenberg et al., 2019).

The other ten mice of the group continued to drink within the IC but they did not initially distinguish between the two different tones. After changing the tone length (from half a second to one second), significant differences in the nosepoke behavior depending on the tone could be detected. The mice did more nosepokes in response to the positive tone compared to the nosepoke number for the negative tone. However, the number of nosepokes for the negative tone increased when the probability of it occurring was increased (up to 50%).

This was particularly evident in the examination of individual learning performance, when nosepoking was barely supressed by the negative tone. Overall, mice made many correct responses for the positive tone, but markedly fewer correct responses for the negative tone. Accordingly, the mice seemed to have a high motivation to perform nosepokes regardless of the outcome.

There was also an increase in the number of visits over the course of the experiment. However, the number of licks per day hardly changed. The explanation might be that the possibility to drink was reduced by increasing the number of trials with the negative tone. Thus, to get the same amount of liquid, more visits had to be made. It may be that the motivation to interact with the nosepoke sensor was so strong that the risk of punishment was accepted. This would be in line with literature data showing that mice continue to operate a lever although it was associated with a stimulation of ‘aversive brain regions’ (Cazala, 1986).

By giving many incorrect responses to the negative tone, the mice also received a correspondingly high number of airpuffs, which in turn could have led to habituation to the airpuff. The punishment would therefore no longer be perceived as a valid punishment (Kahnau et al., 2021). Another explanation could be that the permanent presentation of the tones caused them to no longer be perceived as relevant but rather as a kind of background noise and nosepokes were made independently of the tones.

In conventional tests, mice were placed in a designed test apparatus for a defined test period and are exposed to the stimuli for that defined time (e.g., Bailoo et al., 2018; Boleij et al., 2012; Kloke et al., 2014; Krakenberg et al., 2019; Richter et al., 2012). After the test phase, the mice were transferred back to their home-cages, where they spent their time undisturbed until the next test phase. To the contrary, in our system, which also served as the home-cage, no such breaks occurred. Thus, the stimulus might have had no or little relevance and the focus might be on opening the doors, driven by the motivation to drink.

Our results suggest that rest periods should be included in order to maintain the concentration and/or motivation of the mice. Therefore, for the next developmental step, we decided to schedule breaks, while the mice had access to the water without presentation of the tones, between the individual conditioning and testing phases. To exclude possible influences from previous conditioning phases, we again worked with another naïve mouse group in the next developmental step.

## Developmental Step 3

### Methods

#### Animals

The twelve female mice of group three arrived at the institute in September 2020. At the start of the third developmental step, the mice were six weeks old. At the end of this experiment, the mice were 21 weeks and used in various cognitive experiments (data not published) and in an experiment to develop a home-cage based consumer demand test based on the mouse positioning surveillance system (data not published yet). The mice started barbering behavior at the age of 31 weeks, 10 weeks after the experiment presented here.

#### Corner conditioning protocol

In order to successfully condition the mice of group three to tone-frequencies, further modifications were made to the corner conditioning protocol described earlier. This experiment was pre-registered in Animal Study Registry (animalstudyregistry.org, doi: 10.17590/asr.0000228). In the active corner and after hearing the positive tone-frequency, the IC doors could be opened by a nosepoke for ten seconds. In addition, the tone length as well as the airpuff length was extended to two seconds. The tone-frequencies for the first conditioning phase of group three were the same as for group two (6.814 kHz at 70 dB and 13.629 kHz at 70 dB). For the second conditioning phase, tone-frequencies between 6.814 kHz at 70 dB and 9.636 kHz at 70 dB were used. Also, for group three, the probability of the negative tone was increased step by step (supplementary material table S2).

#### Cognitive bias test

After the conditioning phase, the cognitive bias test followed. This was done by adding ambiguous tone-frequencies, which were calibrated between the positive and negative tone-frequencies (first cognitive bias test: 8.103 kHz, 9.636 kHz, 11.459 Hz, second cognitive bias test: 7.431 kHz, 8.103 kHz, 8.836 kHz). For the determination of these ambiguous tones, geometric mean, which is the perceived middle between two tones, was used. To determine the geometric mean (GM), the square of the product of the two chosen tone frequencies is calculated.

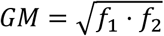

The tritone of the original low and high frequency is then used as the respective high and low frequency to calculate two additional tritones, generating a scale of five tones, each perceptibly equidistant to their neighbors. The SPL was checked with a measuring microphone and the Room Acoustics Software.

The probability for each of the three ambiguous tone frequencies to be played was five percent. By entering the active corner, one of the five different tone-frequencies was randomly presented. The mice received water by performing a nosepoke at the positive tone, and received an airpuff by performing a nosepoke at the negative tone. The mice received neither a reward nor a punishment for the ambiguous tones. For data evaluation, the nosepoke behavior toward the ambiguous tone-frequencies was measured.

During baseline measurement, the housing conditions were as described in section “Home-cage based set-up”. To manipulate the cognitive bias, the housing conditions were changed. The mice had less bedding (2 cm high), less nesting material (four papers), less housing (one mouse house), no running disk, one handling tube, two wooden gnawing sticks, no active enrichment and no resting platform. For further treatment effect, the mice were additionally restrained. For this purpose, the mice were handled by tail and placed in a tube. In the tube, the mice were unable to move and had to remain in the tube for three minutes. This procedure was performed on four consecutive days at 08:00 to 9:30 o’clock during the cognitive bias measurement.

### Results

#### Corner conditioning protocol

From day 57 (figure 10), the tone-frequencies were changed. In total, the mice made more nosepokes in response to the positive tone compared to the number of nosepokes made in response to the negative tone-frequency (*F*_1,429_ = 3578, p < 0.0001). The experimental days also seem to have an influence on the nosepoke number (main effect experimental day: *F*_48,418_ = 4.77, p < 0.0001) as well as the interaction of day and tone (*F*_48,429_ = 6.13, p < 0.0001).

**Figure 10:**
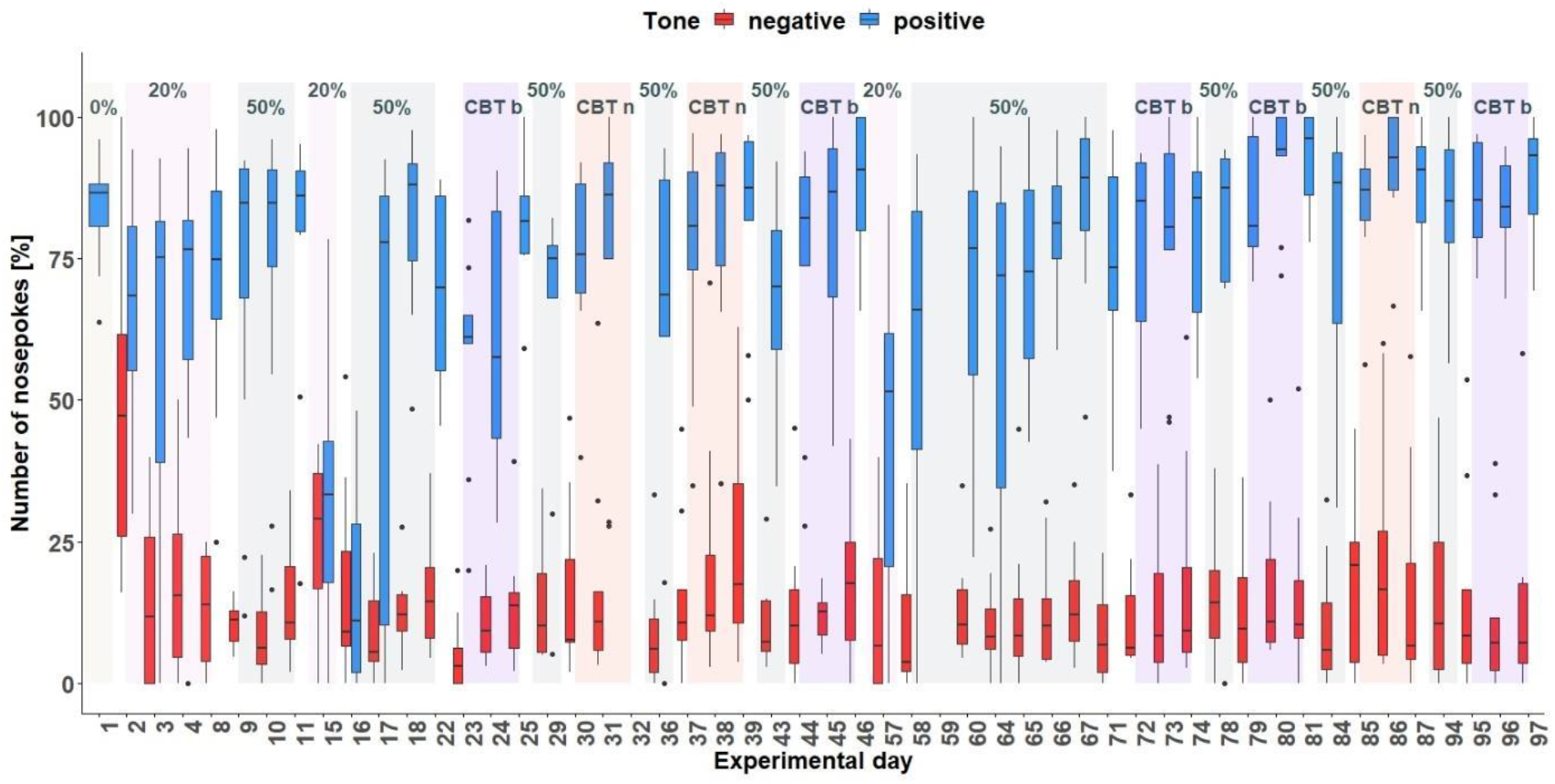
Corner conditioning group three. Number of nosepokes in percent made in response to two tone-frequencies. The data for experimental day 32 and 59 are missing due to technical problems with the IntelliCage system. There was an experimental break between day 47 and 56. After each treatment, no tones were presented. On the y-axis, the number of nosepokes in percent are shown. The x-axis shows the experimental days. During the experimental time period, the tones were presented with different probabilities. 0% = no negative tone, 20% = 20% negative tone probability, 50% = 50% negative tone probability, CBT b = cognitive bias measurement baseline, CBT n = cognitive bias measurement under negative conditions with less bedding and nesting, no enrichment and daily restraining

#### Individual learning success conditioning phase 1

Conditioning phase 1 run for 11 days (figure 11). Experimental day 16 is quite noticeable, where all mice performed worse. It was found that a technical problem occurred during the tone playback. Therefore, for learning success evaluation only 10 days were used.

**Figure 11:**
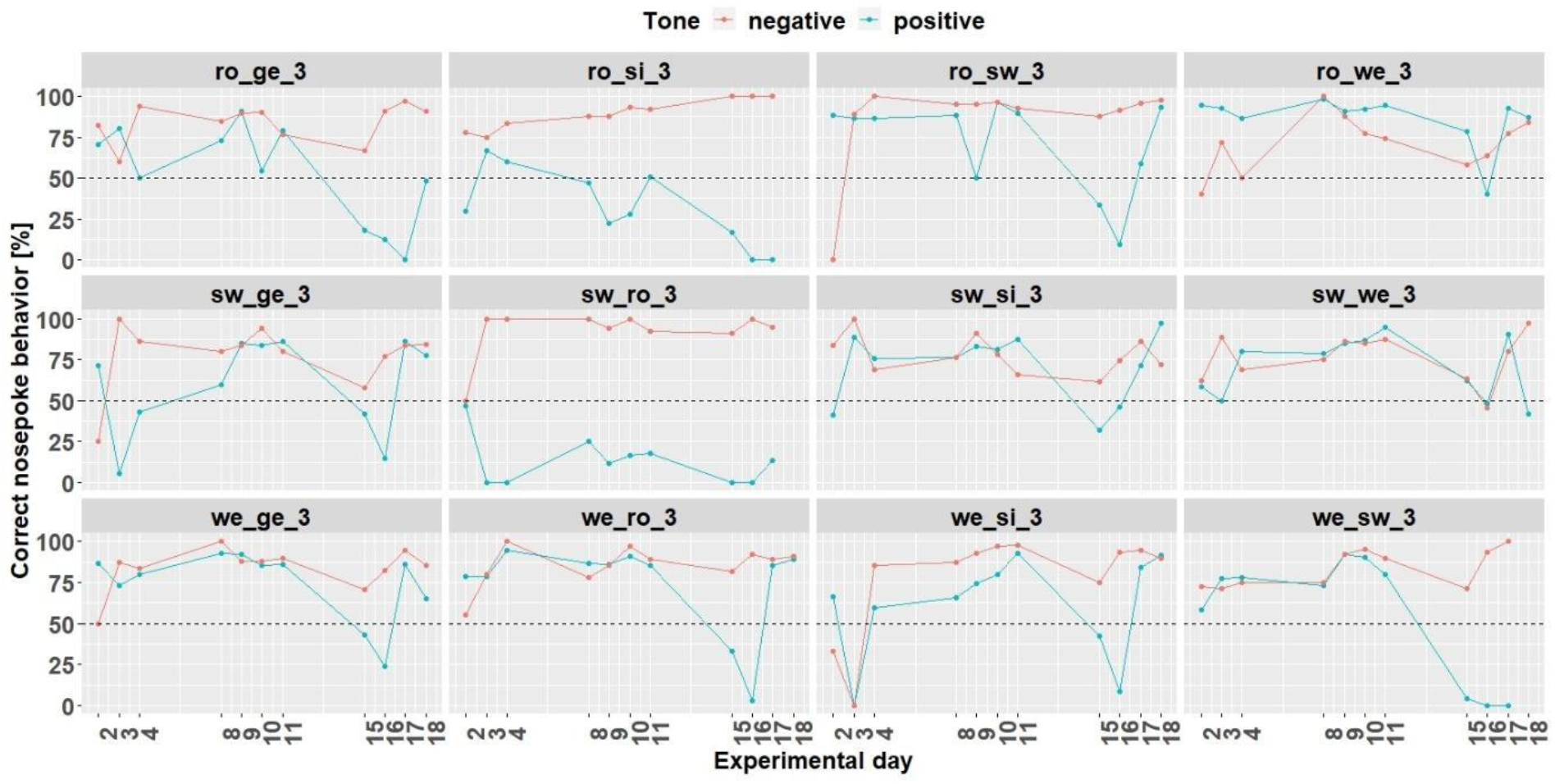
Individual learning success during conditioning phase 1 of mouse group three. On day 16, due to technical problems, the tones were not played correctly The mice ro_si_3, sw_ro_3 and we_sw_3 were excluded from the experiment from day 18 onwards. Learning criterion: 75% of correct nosepoke behavior over 50%.

Nine mice (ro_ge_3, ro_sw_3, ro_we_3, sw_ge_3,sw_si_3, sw_we_3, we_ge_3 we_ro_3 and we_si_3) out of 12 mice reached the learning criterion (75% of correct nosepoke behavior over 50%). The mice ro_si_3, sw_ro_3 and we_sw_3 stopped to drink and were taken out of the experiment at day 18.

#### Cognitive bias test 1

During the first CB test (figure 12), the tone-frequencies influenced the number of nosepokes, which were made after hearing the tone-frequencies (*F*_4,96_ = 28.55, p < 0.0001). A *post hoc* comparison showed except for the negative and near-negative tone (tone-frequency which is close to the negative tone-frequency), the mice discriminated between the different frequencies (table 3). Also the treatment (baseline measurement and negative treatment (less bedding and nesting, no enrichment and daily restraining) has an influence on the nosepoke behavior of the mice (*F*_2,16_ = 5.08, p = 0.02). A *post hoc* comparison showed that the mice made less nosepokes during baseline 1 measurement compared to baseline 2 measurement and negative treatment (table 4). The interaction of tone-frequency and treatment had no influence on the nosepoke behavior (*F*_8,96_= 1.05, p = 0.4).

**Figure 12:**
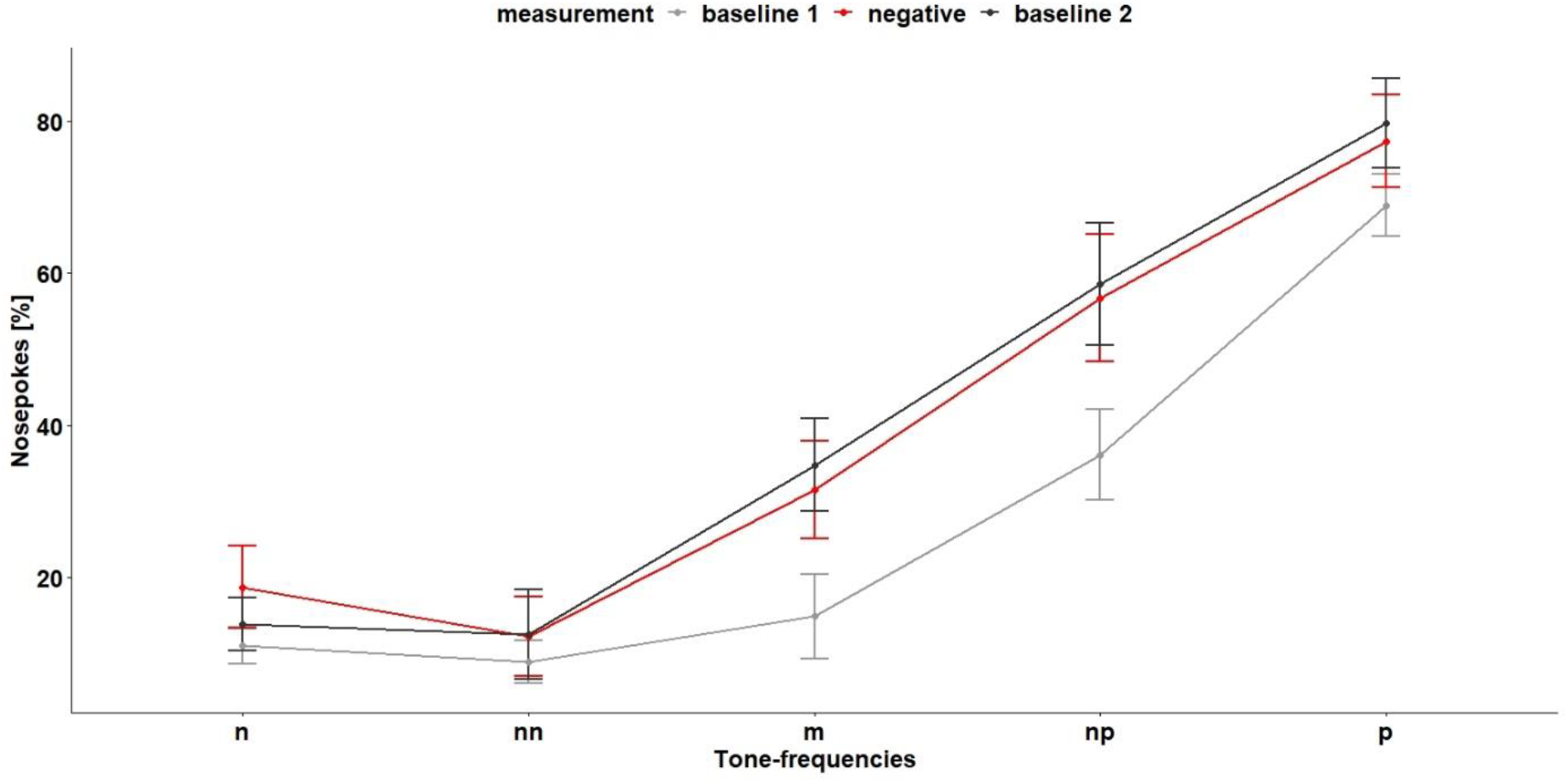
Cognitive bias test 1. The x-axis shows the tone-frequencies with n = negative tone, nn = near-negative tone, m = middle tone, np = near-positive tone and p = positive tone. The y-axis shows the number of nosepokes in percent made in response to the tone-frequencies. During negative measurement the housing conditions were changed compared (less bedding and nesting and no enrichment) to baseline measurement and the mice were restrained daily. n = 9

**Table 3:** Results of the *post hoc* comparison of the performed nosepokes in response to the tone-frequencies for the first cognitive bias test. n = negative tone, nn = near-negative tone, m = middle tone, np = near-positive tone and p = positive tone

| Comparison | Estimate | SE | df | t.Ratio | p-Value |
| --- | --- | --- | --- | --- | --- |
| m – n | 12.52 | 3.87 | 96 | 3.24 | <0.001 |
| m – nn | 15.79 | 3.87 | 96 | 4.08 | <0.001 |
| m – np | -23.41 | 3.87 | 96 | -6.06 | <0.0001 |
| m – p | -48.27 | 3.87 | 96 | -12.48 | <0.0001 |
| n – nn | 3.27 | 3.87 | 96 | 0.85 | 0.4 |
| n – np | -35.93 | 3.87 | 96 | -9.29 | <0.0001 |
| n – p | -60.79 | 3.87 | 96 | -15.72 | <0.0001 |
| nn – np | -39.2 | 3.87 | 96 | -10.14 | <0.0001 |
| nn – p | -64.06 | 3.87 | 96 | -16.57 | <0.0001 |
| np – p | -24.86 | 3.87 | 96 | -6.43 | <0.0001 |

**Table 4:**
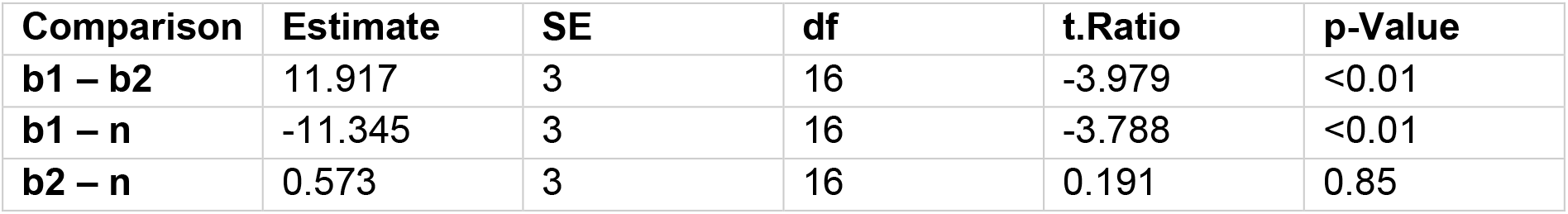
Results of the *post hoc* comparison of the performed nosepokes during baseline measurement and negative treatment in response to the tone-frequencies. b = baseline, n = negative treatment (less bedding and nesting, no enrichment and daily restraining)

#### Individual Learning Success Conditioning Phase 2

Due to technical issues, the data of day 59 are missing and was excluded for learning success evaluation. During conditioning phase 2 (figure 13) 8 (ro_sw_3, ro_we_3, sw_ge_3, sw_si_3, sw_we_3, we_ge_3, we_ro_3 and we_si_3) out of 12 mice reached the learning criterion. The mice ro_si_3 and we_sw_3 stopped to drink and were taken out of the experiment at day 59. The mouse sw_ro_3 stopped to drink, too, and was taken out of the experiment at day 65.

**Figure 13:**
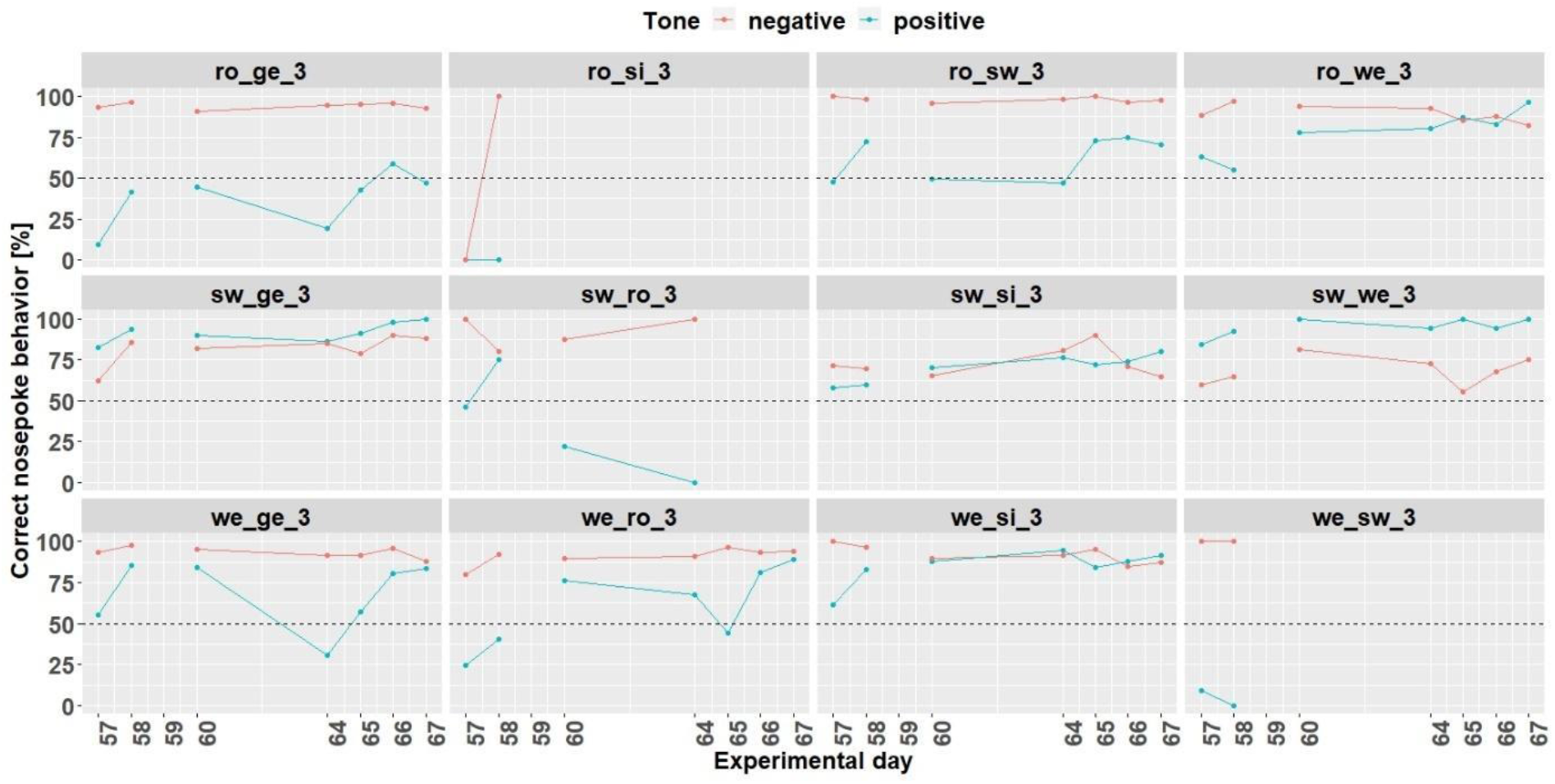
Individual learning success during conditioning phase 2 of mouse group three. Data of day 59 is missing due to technical problems.

Also during the second CB test (figure 14), the tone-frequencies influenced the number of nosepokes (*F*_4,112_ = 27.27, p < 0.0001). Again, the mice did not differentiate between the negative and near-negative tone but between all other tone-frequencies (table 5). The measurement and the interaction of tone-frequency and treatment had no influence on the nosepoke number (main effect treatment: *F*_3,21_ = 1.67, p = 0.2, interaction: *F*_13,112_ = 0.62, p = 0.8).

**Figure 14:**
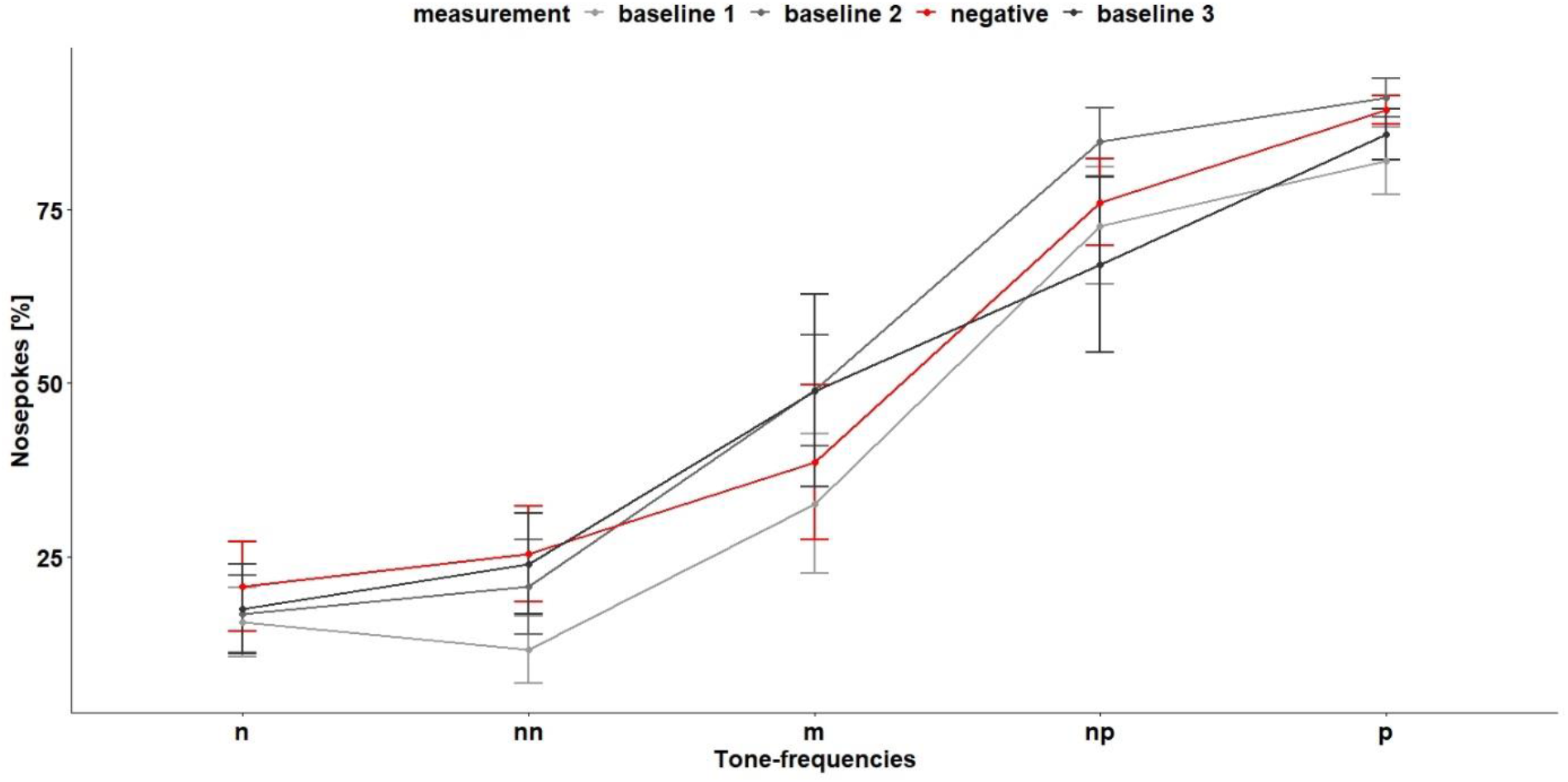
Cognitive bias test 2. The x-axis shows the tone-frequencies with n = negative tone, nn = near-negative tone, m = middle tone, np = near-positive tone and p = positive tone. The y-axis shows the number of nosepokes in percent made in response to the tone-frequencies. During negative treatment the housing conditions were changed compared to baseline measurement and the mice were restrained daily.

**Table 5:** Results of the *post hoc* comparison of the performed nosepokes in response to the tone-frequencies for the first cognitive bias test. n = negative tone, nn = near-negative tone, m = middle tone, np = near-positive tone and p = positive tone

| Comparison | Estimate | SE | df | t.Ratio | p-Value |
| --- | --- | --- | --- | --- | --- |
| m – n | 24.72 | 4.42 | 112 | 5.59 | <0.0001 |
| m – nn | 21.92 | 4.42 | 112 | 5.0 | <0.0001 |
| m – np | -32.89 | 4.42 | 112 | -7.44 | <0.0001 |
| m – p | -44.87 | 4.42 | 112 | -10.15 | <0.0001 |
| n – nn | -2.79 | 4.42 | 112 | -0.63 | 0.53 |
| n – np | -57.61 | 4.42 | 112 | -13.03 | <0.0001 |
| n – p | -69.59 | 4.42 | 112 | -15.74 | <0.0001 |
| nn – np | -54.82 | 4.42 | 112 | -12.4 | <0.0001 |
| nn – p | -66.80 | 4.42 | 112 | -15.1 | <0.0001 |
| np – p | -11.98 | 4.42 | 112 | -2.71 | <0.001 |

#### Body Weight and IntelliCage Behavior

Body weight (figure 15) was influenced by the treatment (*F*_5,963_ = 17.4, p < 0.0001) as well as by the experimental day (*F*_1,963_ = 196, p < 0.0001). Over time, body weight increased continuously. Also the interaction of experimental day and treatment influenced body weight (*F*_5,963_ = 12.52, p < 0.0001). The number of licks (figure 15) over time were influenced by treatment (*F*_5,963_ = 30.79, p < 0.0001) but not by experimental day (*F*_1,963_ = 0.03, p = 0.9) or the interaction of experimental day and treatment (*F* _5,963_= 1.8, p = 0.1). The number of visits (figure 15) were influenced by treatment (*F*_5,963_ = 50.29, p < 0.0001). The analysis showed a tendency towards influence of the experimental day on the visit numbers (*F*_1,963_ = 3.5, p = 0.06). However, the interaction of experimental day and treatment had an influence on the visit number (*F*_5,963_ = 6.8, p < 0.0001).

**Figure 15:**
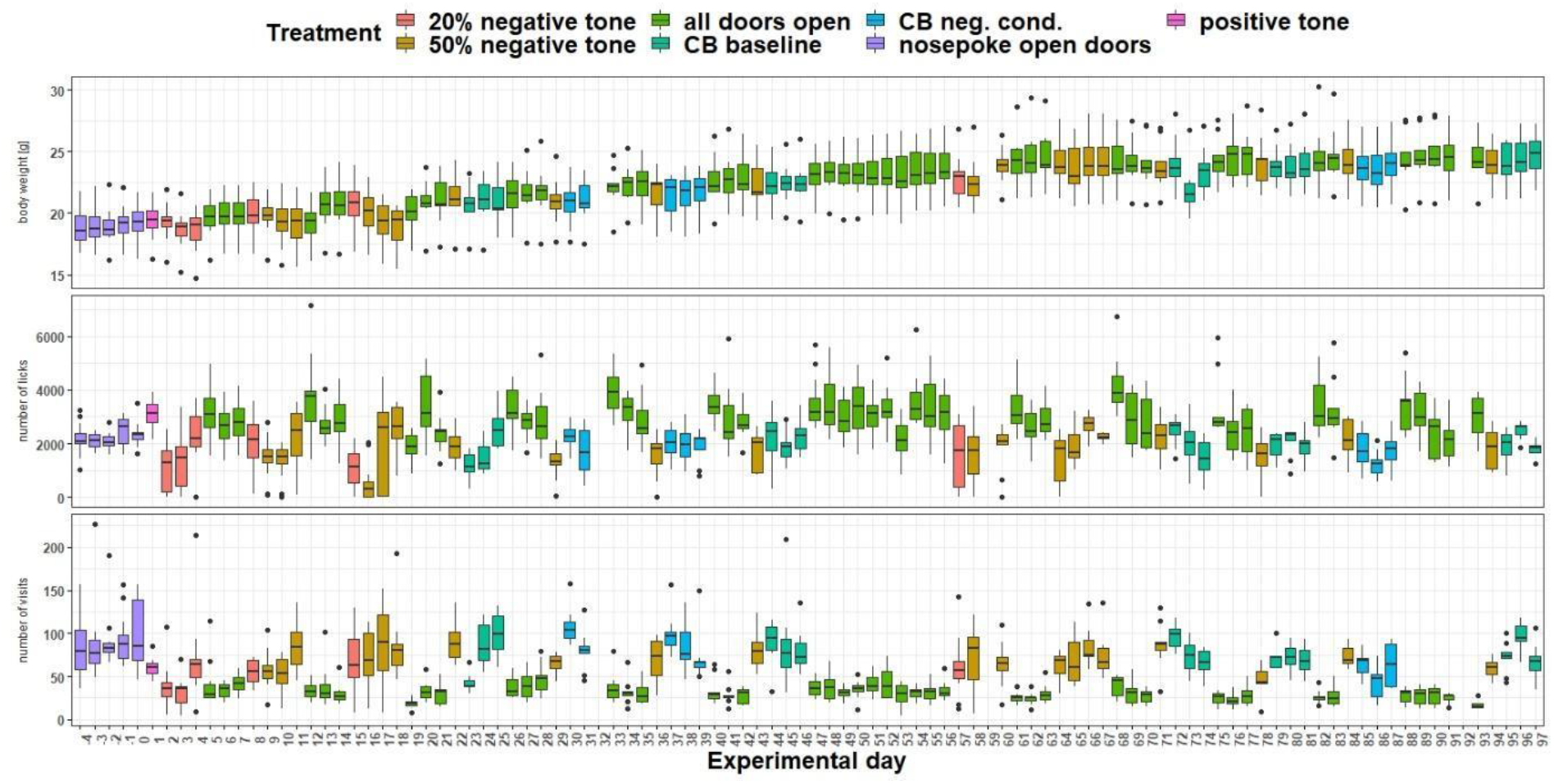
Measurement of body weight, IntelliCage corner visits and lick number over time. The x-axis shows the experimental days. On the y-axis, first the body weight, second the lick number and third the visit number is shown. Different tones with different playback probabilities were presented throughout the experimental period (treatment). The data of experimental day 32, 59 and 92 is missing, due to technical issues.

## Discussion

The third developmental step was also described as ‘corner conditioning protocol’, where tone-frequencies were played whenever a mouse visited the active IC corner. The tone length was changed again (from one to two seconds) compared to developmental step two. The assumption was that this change would allow the mice to discriminate the tone-frequencies more easily. In the study by de Hoz and Nelken, the tone-frequencies were played throughout the complete time of a corner visit. The playing of the tone was stopped only after the mouse left the corner and was re-initiated by a new corner visit (De Hoz & Nelken, 2014). This extreme adjustment of playback length was not considered for our experiment, since it is not known how the individual tone presentation length influences the nosepoke behavior, and thus, the cognitive bias of the mice. There was a potential for individual visit durations to have an influence on the individual mouse assessment of ambiguous tone, making the results difficult to interpret and thus reducing the validity of the data.

Like in developmental step two, some mice in group three could not be conditioned to the tone-frequencies. However, the remaining mice learned effectively and made more nosepokes in response to the positive tone compared to the negative tone. In addition, the mice seemed to be more hesitant in nosepoke behavior compared to the mice in developmental step two. This becomes evident when examining individual learning performance: There were slightly fewer correct responses for the positive tone and more correct responses for the negative tone. This implies that they performed less nosepokes overall, which has a positive effect on the amount of correct answers for the negative tone but a negative effect on the answers for the positive tone. The airpuff seems to be perceived as negative. However, since some mice had to be excluded in this and in the previous developmental step because they stopped drinking, it should be considered whether the airpuff of 0.5 bar is too intense and might be reduced which could reduce the drop-out rate.

In other studies, punishment is not used at all (Graulich et al., 2016; Hintze et al., 2018; Novak et al., 2016; Verjat et al., 2021), as it is discussed that punishment during conditioning and in the test itself may already have an influence on the cognitive bias (Roelofs et al., 2016). However, conditioning with punishment seems to be easier to learn and thus seems to succeed faster (Lagisz et al., 2020). In our system, we chose to use a punishment because the behavior of the mice can be interpreted clearly. The mice want to avoid the airpuff and therefore do not poke when the negative tone is presented. We come to this conclusion based on our experience of mice immediately performing nosepokes upon entering the IC corners if no airpuffs are included in an experimental design.

Lagisz and colleagues identified in their systematic review and meta-analysis that a go/go active choice paradigm (go to receive a reward and go to avoid a punishment) leads to the most sensitive set-up (Lagisz et al., 2020). It is discussed wheather in a go/no-go paradigm the no-go behavior could be related to reduced activity or motivation and not to negative expectation of the future event (Enkel et al., 2010; Matheson et al., 2008). Nevertheless, we chose in our system a go/no-go paradigm. The mice had to nosepoke (go) to receive the reward (water) and not to nosepoke (no-go) to avoid the punishment (airpuff). In addition, the mice had to leave the IC corner and re-enter it (go) to initiate a new trial. We chose a go/no-go paradigm for the same reason that we used the airpuff as a punishment. The behavior in response to the tones is more easily distinguished and interpreted. In addition, by self-initiating the trial, there are no waiting times and the mice have the possibility to complete the trial in a self-determined manner (Hintze et al., 2018; Krakenberg et al., 2019). This choice of experimental design allows us to assume that the mice are highly motivated and facilitates the derivation of a conclusive interpretation of the mice’s behavior.

In the third development step we also analyzed the visit and lick behavior. Both seem to be influenced by the treatment (breaks, conditioning or cognitive bias measurement). By starting conditioning, fewer visits and licks were made. It can be assumed that the lick number is also influenced by the circumstance that the IC doors were permanently open during the breaks. This allowed the mice to drink more per visit during the breaks, which consequently reduced the number of visits and increased the number of licks. The data suggest that the mice need more time to drink, as weight was also affected by the treatment. It would therefore be reasonable to increase IC open-door-time. However, the open-door-time should not be that long that the number of visits is reduced because more licks might be made per visit and thus fewer visits are needed and made overall. This in turn would lead to a reduced number of trials for evaluation.

All three groups of mice showed barbering behavior over the lifespan. This behavior occurred at different ages in the respective groups. Group one showed barbering behavior immediately after the experiments presented here, group two during the experiments and group three a few weeks after the experiments presented here. However, it is likely that the behavior was present earlier, as it was only visible through fur lesions. Barbering is a common behavior in female C57 mice (Garner, 2005; Kahnau et al., 2022B). The reasons for the occurrence of this behavior are still unknown. To gain a better understanding of the behavioral course of barbering, we have developed a score sheet (Kahnau et al., 2022B). Whether and what influence barbering has on the mice and thus on the experimental data is unclear. We assume that the influence on the data presented here is rather low, as we were able to condition mice and measure the cognitive bias. Nevertheless, it is necessary to investigate this behavior further and to report it if it occurs.

Because we assumed successful conditioning in developmental step three, the cognitive bias test followed. With the automated and home-cage based set-up presented here, it was possible to measure the cognitive bias of female C57BL/6J mice. Our data showed a sigmoidal curve of data points decreasing from positive tone-frequency to negative tone-frequency. Our result suggests that the ambiguous tone-frequencies are perceived and interpreted differently with respect to the previously conditioned tone-frequencies, which is a basic requirement of a valid cognitive bias test (Gygax, 2014; Hintze et al., 2018; Krakenberg et al., 2019).

We hypothesized that mice living in enriched housing conditions (from 28 days of age) would be affected in their emotional state by removal of enrichment and additional restraining. In fact, we were able to detect a change in the cognitive bias. The mice showed more nosepoke behavior while kept under negative conditions compared to the time of the first baseline measurement, indicating a positive, optimistic cognitive bias. This increased nosepoke behavior was still evident during the second baseline measurement, when the negative conditions had been eliminated. This result is surprising because studies in rats showed that rats housed under negative housing conditions showed a negative cognitive bias (Burman et al., 2009; Harding et al., 2004) and a transfer from standard to enriched housing conditions led to a shift from pessimistic to optimistic cognitive bias (Brydges et al., 2011; Richter et al., 2012). So far, only Resasco and colleagues were able to measure an influence of housing conditions on cognitive bias in mice. Unlike to our study, enriched housed mice seemed to have a positive expectancy related to the ambiguous stimulus compared to standard housed mice (Resasco et al., 2021).

The question arises why the mice in our experiment seem to have a more optimistic cognitive bias after removing enrichment and with restraining. One explanation might be that the mice experience boredom due to the removal of enrichment, since most stimulating objects have been removed. According to optimal arousal theory, individuals strive for an optimal arousal state. If an individual does not have this arousal state and/or experiences boredom, it would seek something arousing/stimulating. However, if the arousal state is too strong, the individual would seek less arousing stimuli (Mitchell et al., 1984).

In our experiment, this could indicate that the mice do not have an optimal arousal state due to the removal of enrichment and that this is targeted by an increased willingness to take risks to receive an airpuff. However, the mice have also been additionally restrained. Thus, it is not possible to identify which factor (removal of enrichment or restraining) or both factors had an influence on the cognitive bias. The influence also seems to be so strong that an increased nosepoke behavior (compared to the first baseline measurement) could also be detected for the second baseline measurement. This raised the question of whether the mice really had a more optimistic cognitive bias or whether the tones were too “easy” to distinguish. Therefore, we decided to reduce the tone scalar.

The mice also learned to discriminate between tones which were closer to each other, and learned when they received water and when they received an airpuff. Therefore, another cognitive bias test was performed.

During the second test phase, we again observed a sigmoidal curve in the data, but no change in cognitive bias. This result is consistent with other studies (Bailoo et al., 2018; Bračić et al., 2022). It should be noted that during the second test phase, the period of negative conditions was significantly lower. It is feasible that one week has no influence on the cognitive bias of mice or that the experiences already made have led to a kind of habituation. It is also possible that the test systems developed so far, including the system presented here, are not sensitive enough. In addition, the possible change in cognitive bias might not last long enough to be measured or is covered by positive stimulation due to cognition training (Krakenberg et al., 2019). Another reason why we could not measure a change could be that a group of mice serving as their own control is not informative enough, as we cannot rule out a temporal carrying over effect for the second baseline. However, the study of Bracic and colleagues showed that the cognitive bias was repeatable over multiple measurements (Bračić et al., 2022). Further experiments are necessary to better interpret the results presented here. For example, it is necessary to test whether a mouse group can serve as its own control group.

It should be noted that too frequent repetition of presenting the ambiguous stimuli could also lead to mice learning that neither reward nor punishment occurs with ambiguous stimuli (Roelofs et al., 2016). Therefore, it is necessary to ensure that the ambiguous stimuli are distributed in an appropriately high trial number of positive and negative stimuli (Krakenberg et al., 2019).

In some mouse test systems, the trial number per session is relatively low, i.e.1-32 trials (Boleij et al., 2012; Kloke et al., 2014; Novak et al., 2016). In contrast, in the set-up of Hintze and colleagues and in the automated touch-screen system of Krakenberg and colleagues, up to 54 trials per day could be performed. However, it was always necessary to remove the mice from their familiar environment, thus separating them from their group members (with the exception of individual housing) and determining the time of the test, which could have an influence on the motivation to participate in the test.

In our set-up, the number of trials varied depending on how frequently the IC corners were visited (group three: 4 - 214 visits = trials), but were distributed over the entire day. The animals decided independently from the experimenter when to enter the IC (if the IC was not already occupied by another mouse), which makes a high motivation to participate in the test plausible. Even though only one mouse could be in the IC at a time, it was possible for all mice to enter the IC several times a day, and thus, initiate trials in the IC itself. This was also shown in an automated and home-cage based consumer demand test, for which a similar test setup was used as described here (Kahnau et al., 2022A). It is also not necessary to manipulate the night/day rhythm (as e.g., in Krakenberg et al., 2019), as in experiments in which the presence of an experimenter during data acquisition is required. This, in turn drastically reduces the time required (daily control of animals and set-up of about 30 minutes).

## Conclusion

The cognitive bias test seems to be a suitable test method to measure the affective state of animals (Lagisz et al., 2020). So far, however, these tests are very labor intensive and require animals to be tested outside of their home cages, which has implications for the animals and thus the data. (e.g., Bračić et al., 2022; Hintze et al., 2018; Kloke et al., 2014; Krakenberg et al., 2019). Therefore, we aimed to develop an automated and home-cage based cognitive bias test for mice.

In the study presented here, we describe the developmental steps for such a test concluding in a method that allows measuring the cognitive bias in mice. By presenting the various stages of development, we intended to provide a better understanding of the structure of the test method. We also contribute to providing comprehensive information to the scientific community that can be used to develop further automated and home-cage based systems.

Automation and home-cage based testing offers the advantages of testing the mice in their familiar environment and during their active phase. The influence of the animals on each other is reduced, as only one mouse can be in the test-cage at a time. Also the influence of the experimenter is reduced to a minimum. The fact that the mice can choose the time of the experiment and initiate trials themselves gives them control over what occurs and suggests that the mice are highly motivated. All this, in turn, might have a positive impact on the validity of the data.

We were able to measure and manipulate the cognitive bias of the mice although further research is needed for a better understanding of the mice’s cognitive bias. We will continue to develop our test system and use it to assess the burden of commonly used behavioral tests such as the Water Maze Test, and to include the perspective of the mouse in this assessment.

## Supplementary Information

The supplementary material and raw data are online available at: https://zenodo.org/record/7224991#.Y0_d1vzP1PY

## Ethical Approval

All experiments were approved by the Berlin state authority, Landesamt für Gesundheit und Soziales (LAGeSo), under license No. G 0182/17 and were in accordance with the German Animal Protection Law (TierSchG, TierSchVersV).

## Authors Contribution

Conceptualization: Pia Kahnau, Lars Lewejohann, Anne Jaap

Methodology: Pia Kahnau

Technical support: Birk Urmersbach

Analysis: Pia Kahnau

Writing – original draft preparation: Pia Kahnau

Writing – review and editing: Pia Kahnau, Anne Jaap, Birk Urmersbach, Kai Diederich, Lars Lewejohann

Funding acquisition: Lars Lewejohann

Supervision: Lars Lewejohann

## Acknowledgements

The authors thank the animal caretakers of the German Federal Institute for Risk Assessment, especially Carola Schwarck, Lisa Gordijenko and Heiderose Camus, for their support in the animal husbandry.

